# High quality chromosomal genome assemblies of three human Plasmodium species directly from natural infections

**DOI:** 10.64898/2026.02.16.706136

**Authors:** Sunil Kumar Dogga, Jesse C. Rop, Alex Makunin, Fiona Teltscher, Damon-Lee Pointon, Ying Sims, Marcela Uliano-Silva, James Torrance, Thomas C. Mathers, Jonathan M.D. Wood, Sekou Sissoko, Antoine Dara, Dinkorma T. Ouologuem, Arthur M. Talman, Abdoulaye A. Djimdé, Mara K. N. Lawniczak

## Abstract

Malaria, caused by mosquito-transmitted *Plasmodium* parasites, remains a major global health challenge. While *P. falciparum* accounts for most malaria deaths, the lesser-studied *P. ovale* spp. and *P. malariae* also contribute to disease burden across many endemic regions, typically causing milder but chronic infections. Here, we generated high-quality, chromosome-level reference genomes for *P. ovale wallikeri*, *P. malariae,* and *P. falciparum* directly from blood samples taken from natural infections. Using PacBio Ultra-Low-Input HiFi long-read sequencing and Hi-C chromatin conformation capture, we overcome limitations posed by low parasite biomass and the absence of *in vitro* culture for *P. ovale wallikeri* and *P. malariae.* These high quality reference genomes resolve previously inaccessible subtelomeric regions that include expanded gene families likely involved in host-parasite interactions, and they enhance our understanding of parasite persistence, transmission, immune evasion and comparative evolutionary studies. This work provides a critical foundation for including these neglected species in global malaria elimination efforts.

## Main

Malaria remains a major global health issue, with half of the world’s population thought to be at risk (WHO). In humans, the disease is caused by *Plasmodium falciparum*, *P. vivax*, *P. knowlesi*, *P. ovale*, and *P. malariae* species. While *P. falciparum* continues to be the primary cause of morbidity and mortality in sub-Saharan Africa (SSA), non-*falciparum* species such as *P. ovale* and *P. malariae* are increasingly recognized as contributors to the malaria burden. These species were often overlooked due to misdiagnosis, limited molecular surveillance, and a lack of dedicated research focus ^1,2^. *P. ovale* is also understood to be two genetically distinct but morphologically identical species, *P. ovale curtisi* and *P. ovale wallikeri,* which are sympatric, non-recombining, and differ subtly in latency periods ^3,4^. Across West Africa, where both *P. ovale* species circulate, the relative contribution of each remains poorly characterized due to the absence of accurate diagnostic markers. These parasites are believed to form hypnozoites in the liver, similar to *P. vivax*, allowing them to cause relapsing infections long after the initial exposure ^5^, although the evidence for this is minimal ^6^. *P. malariae*, on the other hand, is capable of establishing chronic blood infections that may persist asymptomatically for years or even decades ^7^. Although traditionally considered to cause relatively mild illness, *P. ovale* and *P. malariae* have both been associated with serious complications, including severe anemia, splenomegaly, and in rare cases, congenital malaria ^8–12^. *P. malariae* has also been specifically implicated in nephrotic syndrome ^13,14^. Morbidity associated with long-term carriage of these malaria species is understudied and these infections are likely underreported, in part due to diagnostic limitations and frequent co-infections with *P. falciparum*, which obscures clinical attribution. Recent reports suggest that over half of *P. ovale* and *P. malariae* infections in SSA occur as mixed infections with *P. falciparum* ^1,2,15^.

The prevalence of *P. ovale* spp. and *P. malariae* has been shown to be stable even in areas where *P. falciparum* incidence is diminishing ^16–18^, marking these species as a distinct challenge in eradication efforts. Despite this growing epidemiological relevance, the biology and genetics of *P. ovale* spp. And *P. malariae* remain insufficiently studied due to the absence of continuous *in vitro* culture systems and challenges in accessing clinical samples due to the their typical occurrence as mixed infections and the difficulty of accurate species diagnosis in field settings.

From a global health perspective, the neglect of these parasites poses several risks. They were increasingly identified in travel-related malaria cases, often evading standard prophylaxis, which highlights their potential for silent spread and reintroduction into previously malaria-free regions ^19–21^. Moreover, reports of resistance-conferring mutations, such as those in the *dhfr* gene, raise concerns about the future efficacy of current antimalarial regimens ^22,23^, especially when these parasites are often co-exposed to *P. falciparum* therapeutic regimen which may be suboptimal and provide good resistance-selecting conditions.

The molecular toolkits available to study these species have only recently begun to expand. For *P. falciparum*, the 3D7 reference genome was first assembled over two decades ago ^24^ and later improved using PacBio long-read sequencing ^25^. The nearly telomere-to-telomere representation of all 14 chromosomes resolves most centromeric and subtelomeric regions and remains a highly curated, gold-standard reference, albeit from a laboratory-adapted strain. On the other hand, for *P. ovale* spp. and *P. malariae*, the first reference assemblies (PocGH01, PowCR01, and PmUG01) were published in 2017 ^26^. The *P. malariae* PmUG01 assembly was generated using PacBio long reads and iterative polishing, markedly improving contiguity (61 scaffolds with 14 gapless chromosome contigs, scaffold N50 ≈ 2.3 Mb), whereas the *P. ovale curtisi* and *wallikeri* assemblies in this study were derived largely from short-read Illumina data. Subsequent updates have incorporated long-read sequencing and selective whole-genome amplification (sWGA) to enhance contiguity and gene annotation, particularly in subtelomeric multigene families ^27–29^. The recent *P. ovale curtisi* and *wallikeri* reference genomes (Poc221 and Pow222) generated from clinical samples obtained from travellers returning to the UK, were produced using a hybrid approach that sequenced both sWGA-enriched and non-enriched material with Oxford Nanopore Technology (ONT) MinION under adaptive sampling conditions to deplete human DNA ^28^. The ONT reads had mean lengths of 1.15 Kbp and 1.17 Kbp for Poc221 and Pow222, respectively. The resulting assemblies (scaffold N50s of 217 Kb for Poc221 and 270 Kb for Pow222) represent improvements in contiguity and gene annotation, particularly across subtelomeric multigene families such as *pir*, *phist*, and *resa*-like genes. However, sWGA introduces uneven coverage across the genome ^30^, resulting in fragmented assemblies and incomplete or inaccurate gene annotations, evidenced by the high number of scaffolds (1177 and 787) and the large fraction of genome present as unlocalised contigs (∼35%). Also, subtelomeric multigene families were often poorly covered by sWGA primer sets and may therefore be underrepresented in sWGA-derived data. Moreover, the absence of Hi-C or other chromatin-conformation data for *P. ovale* spp. and *P. malariae* has prevented chromosome-scale scaffolding, leaving numerous contigs unlocalised and not associated with chromosomes and structural breaks across all published references. Collectively, these factors continue to limit comprehensive analyses of genome architecture, synteny, and subtelomeric gene family diversity. Overall, the best reference genomes currently available are the sWGA-enriched ONT assemblies Poc221 and Pow222 for *P. ovale curtisi* and *P. ovale wallikeri*, respectively, and the PacBio long-read based *P. malariae* PmUG01.

Here, we present high-quality, chromosome-level genome assemblies of *P. ovale wallikeri*, *P. malariae*, and *P. falciparum*, generated from clinical infections in Faladie, Mali, a hyperendemic region characterized by intense seasonal malaria transmission. After the elimination of leukocytes and uninfected RBCs, we obtained enriched parasite material from clinical infections, generating long read PacBio data from as little as 2 ng of DNA without culture adaptation or sWGA. Using the Ultra-Low Input (ULI) PacBio long-read sequencing kit and Hi-C, we reconstructed near-complete genomes with greatly improved contiguity and annotation, particularly in subtelomeric regions enriched for multigene families. These assemblies provide an essential foundation for understanding the biology, gene function, and evolutionary relationships of these neglected malaria parasites.

## Results

### Processing, sequencing, and curation of genome assemblies from natural malaria infections

To generate high quality genome references, we enriched parasite fractions from the blood samples of eighteen study participants in Faladie (Mali), representing both symptomatic and asymptomatic malaria cases with varying mixed species infection status. Eight of these samples yielded sufficient parasite material to generate high-quality reference genome assemblies (**Table 1, Extended Data Fig. 1a**). Intravenous whole blood was processed without further culturing using a previously published protocol that enriches parasites using Magnetic-activated cell sorting (MACS) and Streptolysin-O (SLO) (**Extended Data Fig. 1a)** ^31^. High Molecular Weight DNA was then sequenced with PacBio’s Ultra Low Input High-Fidelity (ULI-HiFi) long-read sequencing protocol. Mapping HiFi reads to the *P. ovale* spp. reference genomes (PocGH01, https://plasmodb.org/: v68 and Pow222, ^28^), and examining the *cox1* locus confirmed all *P. ovale* spp. isolates as *P. ovale wallikeri* (**Extended Data Fig. 1b**).

**Table 1:** Assembly summary for all the specimens, in comparison to current published references. The highest-quality assemblies for each species are indicated in green. Specimens used for generating Hi-C data are indicated with a * next to the participant ID, and # is used to indicate those with mixed infections.

|  | Donor ID | ID<br>(Source) | toID | Full assembly |  |  |  |  | Main Chromosomes |  |  |  | chrM<br>(bp) | chrA<br>(bp) |
| --- | --- | --- | --- | --- | --- | --- | --- | --- | --- | --- | --- | --- | --- | --- |
|  |  |  |  | n | Size<br>(Mbp) | Gain<br>(Mbp) | GC<br>% | Gaps | Size<br>(Mbp) | Gain<br>(Mbp) | GC<br>% | Gaps |  |  |
| *P. ovale wallikeri*, Pow | * MSC23 | PowML03 | pxPlaOval3 | 15 | 38.05 | 3.66 | 28.37 | 36 | 37.83 | 12.01 | 28.42 | 9 | 5975 | 34363 |
|  | MSC32 | PowML04 | pxPlaOval4 | 17 | 34.26 | -0.13 | 28.85 | 32 | 32.03 | 6.21 | 29.38 | 32 | 5980 | 29156 |
|  | MSC31 | PowML08 | pxPlaOval8 | 83 | 34.90 | 0.51 | 28.99 | 166 | 29.53 | 3.71 | 30.38 | 165 | --- | 32892 |
|  | MSC25 | PowML09 | pxPlaOval9 | 24 | 37.99 | 3.60 | 28.36 | 58 | 34.95 | 9.13 | 28.99 | 58 | 5970 | 33620 |
|  | --- | Pow222<br>(Nigeria) | --- | 787 | 34.39 | --- | 29.13 | 182 | 25.82 | --- | 31.58 | 182 | 5975 | 34310 |
|  | --- | PowCR01<br>(Ghana) | --- | 779 | 33.53 | -0.86 | 29.26 | 1255 | 20.92 | -4.90 | 33.14 | 1167 | 5584 | 31160 |
| *P. malariae*, Pm | * MSC29 # | PmML03 | pxPlaMala3 | 14 | 36.78 | 3.20 | 23.93 | 13 | 36.78 | 7.24 | 23.93 | 9 | 5966 | 35642 |
|  | MSC19 | PmML04 | pxPlaMala4 | 17 | 35.20 | 1.62 | 24.16 | 18 | 34.94 | 5.40 | 24.20 | 18 | 5967 | 33199 |
|  | MSC28 # | PmML07 | pxPlaMala7 | 75 | 26.08 | -7.50 | 25.05 | 422 | 22.64 | -6.90 | 25.98 | 399 | --- | --- |
|  | --- | PmUG01<br>(Uganda) | --- | 61 | 33.58 | --- | 24.39 | 0 | 29.54 | --- | 25.12 | 0 | 5969 | 34324 |
|  | *P. Brasilianum*<br>(Bolivian I) |  | --- | 43 | 31.37 | --- | 24.81 | 8 | 29.08 | --- | 25.16 | 7 | 6302 | 28948 |
| *P. falciparum*, Pf | MSC28 # | PfML46 | pxPlaFalc46 | 34 | 23.88 | 0.59 | 19.61 | 24 | 23.21 | -0.08 | 19.32 | 23 | 5967 | 32080 |
|  | MSC61 | PfML47 | pxPlaFalc47 | 16 | 23.37 | 0.08 | 19.40 | 7 | 23.30 | 0.01 | 19.37 | 7 | 5966 | 34244 |
|  | MSC29 # | PfML72 | pxPlaFalc72 | 14 | 23.21 | -0.08 | 19.35 | 31 | 23.21 | -0.08 | 19.35 | 31 | --- | 32282 |
|  | --- | Pf3D7 | --- | 14 | 23.29 | --- | 19.34 | 0 | 23.29 | --- | 19.34 | 0 | 5967 | 34250 |

Two Arima Hi-C libraries were generated and sequenced on Illumina platforms, one from a participant with a *P. ovale wallikeri* infection (MSC23) and one from a participant with a mixed *P. malariae*/*P. falciparum* infection (MSC29). These datasets were used to scaffold PacBio ULI contigs (**Fig. 1, Extended Data Fig. 1a,c-e, Methods**). The ULI approach enabled high-quality genome assemblies from specimens with low DNA quantities for *P. ovale wallikeri* (MSC23, 25, 31, 32), *P. malariae* (MSC 19, 28, 29), and *P. falciparum* (MSC28, 29, 61) (**Extended Data Fig. 1a)**. PacBio HiFi reads were depleted of host sequences by mapping to the GRCh38 (GENCODE v32/Ensembl 98) human reference genome, retaining only unmapped reads for assembly with hifiasm (v0.16) ^32^ (**Supplementary Table 1)**. MSC28 and MSC29 were from *P. falciparum*/*P. malariae* mixed species infections, enabling recovery of assemblies for both parasite species, resulting in ten *Plasmodium* genomes from the eight participants.

**Fig. 1.**
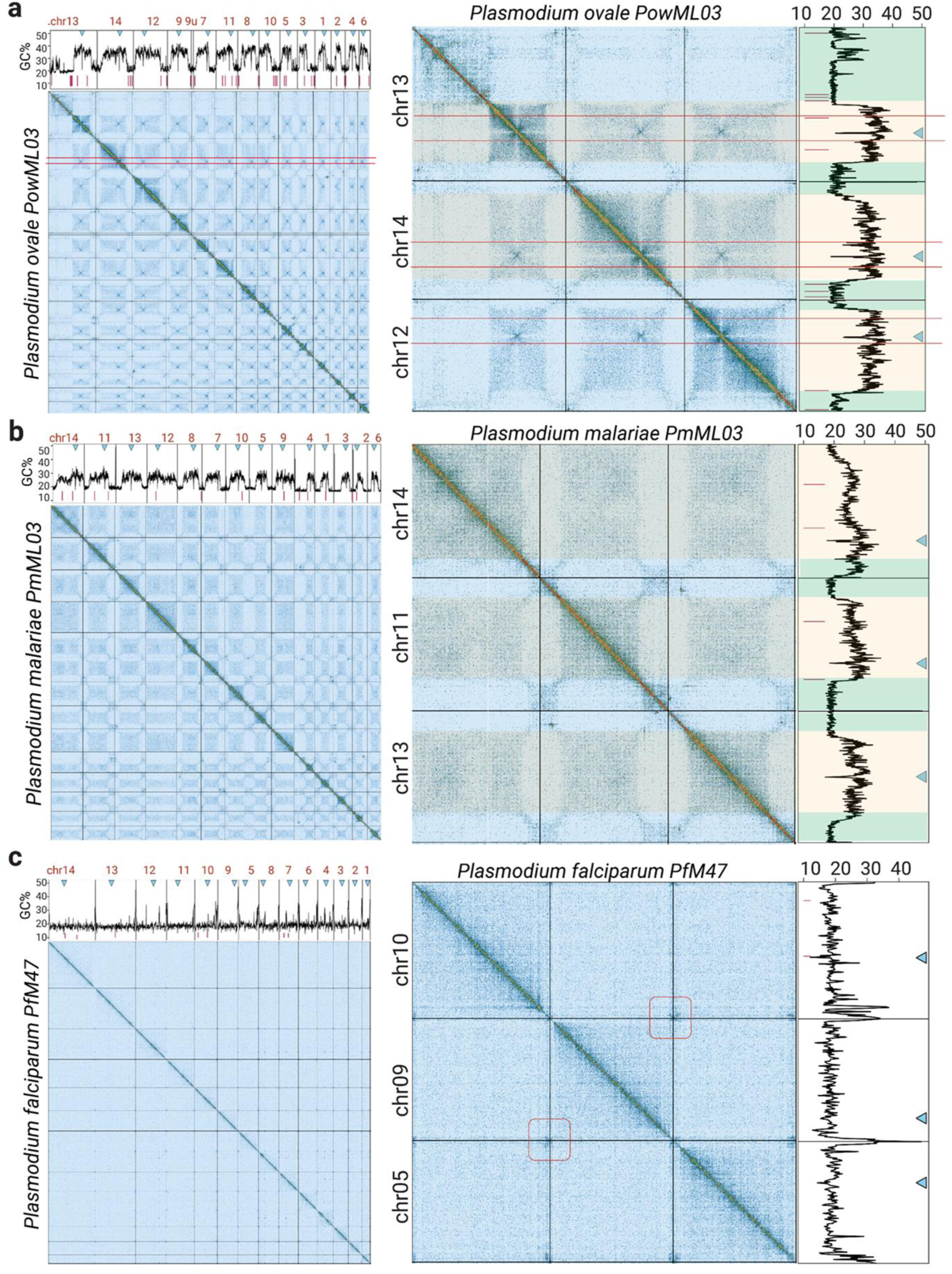
a-c, (left panels) Pretext file snapshot of Hi-C contact maps of super scaffolds (chromosomes) of **(a)** PowML03, (**b**) PmML03, and (**c**) PfML47, arranged by size. GC content was calculated in 10,000 bp non-overlapping bins and plotted along each chromosome, with vertical black lines indicating chromosome boundaries and red segments under the GC lineplot corresponding to breaks in the assembly. (Right panels) The GC content profile mirrors chromosome-scale patterns observed in the Hi-C contact maps, with a high-GC core in central regions of chromosomes (yellow) and reduced GC content in the subtelomeric regions (green) for PowML03 and PmML03. Centromere coordinates, listed in **Supplementary Table 2**, of PowML03 and PmML03 were determined using the cross-shaped interaction patterns typically associated with centromeric domains, indicated by blue inverted triangles. GC content was used to infer centromeric regions in *P. falciparum*, by selecting loci where GC content dropped below 5% across two consecutive 1000 bp bins. While interchromosomal interactions are not apparent in PfML47, indications of interactions at the subtelomeric regions of PfML47 are highlighted with maroon boxes in **(c).**

Genome contiguity was assessed via N50 statistics and comparison with existing references (**Fig. 2a-c, Extended Data Fig. 2a, Supplementary Table 1, Table 1**). Initial hifiasm assemblies ranged from 51-605 contigs, with contig N50 values from 149,862 bp to 1,839,254 bp, the latter being nearly two-thirds the size of the largest chromosome of *P. falciparum* 3D7 (3.29 Mbp) and larger than the theoretical maximum N50 for *P. falciparum* 3D7, which is ≈1.688 Mb, corresponding to chromosome 10, indicating high assembly contiguity (**Supplementary Table 1**). Hi-C data from MSC23 (a *P. ovale wallikeri* infection) was used to scaffold all *P. ovale wallikeri* assemblies, and that from MSC29 (participant with *P. falciparum/P. malariae* mixed infection) was used to scaffold all *P. malariae* and *P. falciparum* assemblies with the tool YaHS ^33^, followed by manual curation. This approach enabled the scaffolding of the hifiasm contigs from the *P. ovale wallikeri* assembly (PowML03), *P. malariae* assembly (PmML03) and *P. falciparum* assembly (PfML47) into the 14 chromosomes of the final assemblies, which together represented the highest-quality assemblies for the three species, with improvements in genome size, contiguity, and chromosomal completeness (**Fig. 2a-c**). A single contig of PowML03, that likely belongs to chr09 based on Hi-C contact maps, remained unplaced and labelled chr09u. PowML03 and PmML03 were each scaffolded using Hi-C data derived from their respective specimens, while PfML47 was scaffolded using Hi-C data from another participant, MSC29 (which yielded the PmML03 and PfML72 assemblies) (**Extended Data Fig. 1a**). Mitochondrial and apicoplast genomes were identified from the contigs using MitoHiFi (**Table 1**) ^34^.

**Fig. 2.**
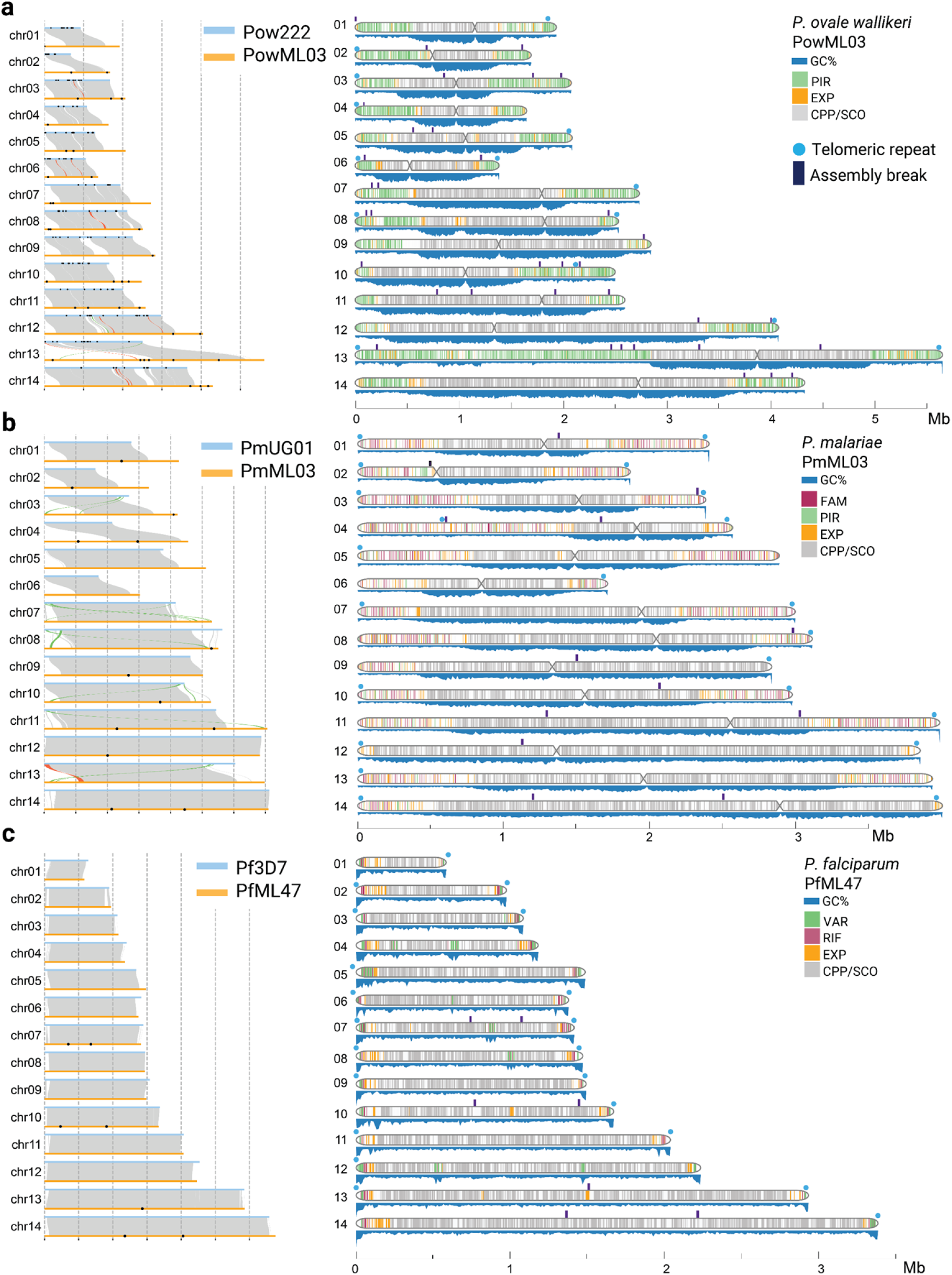
**a-c**, (left panels) SyRI synteny plot of each species’ best assembly for **(a)** *P. ovale wallikeri* (PowML03), **(b)** *P. malariae* (PmML03) and **(c)** *P. falciparum* (PfML47) in comparison to the existing best references, Pow222, PmUG01 and Pf3D7, respectively. (right panels) Chromosome plots for (**a)** *P. ovale wallikeri* (PowML03), (**b**) *P. malariae* (PmML03), and (**c**) *P. falciparum* (PfML47) showing the GC content (blue) in 10kbp bins scaled from a minimum of 15%, presence of genes coding for multigene families (in color) and single copy orthologs (SCO) and/or conserved *Plasmodium* proteins (CPP) (grey), across the 14 chromosomes. Breaks in the assemblies (black tick marks) and presence of telomeric repeats (blue circles) are displayed above the chromosome ideograms. Regions of low GC content in PowML03 and PmML03 correspond to multigene family clusters that overlap with domains observed in the Hi-C contact maps.

### Refined reference assemblies with improved contiguity for *P. ovale wallikeri, P. malariae* and *P. falciparum*

Colinearity with published *Plasmodium* genomes was assessed with genome alignment dotplots, using D-Genies ^35^ which allowed us to orient the chromosomes and manually assign IDs to each assembled chromosome (**Extended Data Fig. 2a-c**). In addition, pairwise whole genome comparisons with best existing references were performed using SyRI ^36^ revealing extended subtelomeric regions in the new assemblies (**Fig. 2a-c, Extended Data Fig. 1g**). Coverage analysis revealed relatively uniform coverage depth across the chromosomes, with <1% of 10 kbp bins covered by < 25 reads (16, 33 and 0 bins respectively for PowML03, PmML03 and PfML47) (**Extended Data Fig. 2d-f**). Mitochondrial copy number was estimated as the ratio of median coverage of mitochondrial to nuclear genomes: the resulting values ranged from 12–86 in *P. ovale wallikeri*, 13–84 in *P. malariae*, and 24–637 in *P. falciparum*, likely reflecting differences in stage composition and associated mitochondrial copy number of the samples (**Extended Data Fig. 2f,g**).

In comparison to published references (PowCR01, Pow222, PmUG01, Pf3D7), we noticed a considerable gain in genome assembly span across both *P. malariae* and *P. ovale wallikeri* **(Table 1, Fig. 2a-c, Extended Data Fig. 2a-c**). For both these species, a majority of the contiguity gains were in sub-telomeric regions (**Fig. 2a-c, Table 1, Extended Data Fig. 2a-c**). For *P. ovale wallikeri*, we saw a reduction in the number of gaps from 1167 (PowCR01) and 182 (Pow222) to 36 (PowML03). With PmML03, 13 gaps remain, compared to 0 gaps in PmUG01 (annotated on **Fig. 1** and **Fig. 2**). We observed a good contiguity concordance of PfML47 with the *P. falciparum* 3D7 reference and only 7 gaps remaining (vs 0 for 3D7). Chromosome lengths were extended for all chromosomes of PowML03 when compared to Pow222, and 11 of the 14 chromosomes of PmML03 when compared to PmUG01 (**Extended Data Fig. 2h**), with the biggest increase seen for chr13 (+ 3,100,788 bp) for PowML03 and for chr04 (+ 1,197,753 bp) for PmML03. Further illustrating the completeness of the chromosomes, repeat sequences (GGGTT[T/C]A) ^37^ associated with *Plasmodium* telomeres were identified at chromosomal ends in a majority of the curated chromosomes, with 14, 22 and 21 loci (of 28 expected) present in chromosomes for PowML03, PmML03 and PfML47, respectively (**Fig. 2a-c, Extended Data Fig. 2i**). Altogether, PowML03 and PmML03 constitute the most complete genomes to date for *P. ovale wallikeri* and *P. malariae*, respectively and PfML46, PfML47 and PfML72 represent high-contiguity, chromosome-level assemblies for uncultured *P. falciparum* isolates from Mali.

### Distinct chromosomal architecture and centromere clustering patterns among *P. ovale wallikeri*, *P. malariae*, and *P. falciparum*

Hi-C contact maps used for scaffolding the assemblies for *P. ovale wallikeri*, *P. malariae* and *P. falciparum* exhibit strong chromosome-scale diagonals with internal substructure (**Fig. 1a-c**, **Extended Data Fig. 1c-e**). The centromere locations in PowML03 and PmML03 were pinpointed using Hi-C contact maps visualised in PretextView, and displayed the characteristic cross-shaped interaction pattern coupled with negative signal indicating reduced interaction with surrounding regions, associated with centromeric domains (**Fig. 1a-c**, **Extended Data Fig. 1c-e, Supplementary Table 2**). In PowML03, we see x-like hotspots along all the chromosomes align with the centromere locations (**Fig. 1a, example within maroon colored lines**). For higher eukaryotes, similar signals can be driven by repetitive nature of centromeres, but in *Plasmodium*, no repeats are consistently associated with cross-like patterns in putative centromeres ^38,39^. This indicates that the cross-like patterns are likely driven by interchromosomal interactions and strong centromere clustering in *P. ovale wallikeri*, and not sequence similarity. This is consistent with previous Hi-C and microscopy studies in *P. falciparum, P. berghei*, *Toxoplasma gondii and Babesia microti* where centromeres co-localize into a clustered nuclear focus ^40–42^. Such clustering is less pronounced in *P. malariae* (PmML03) and *P. falciparum* (PfML47), possibly reflecting weaker Hi-C signal or more dispersed centromere organization in the stages profiled (**Fig 1b,c**). The reduced Hi-C signal observed in PmML03 and PfML47 may reflect differences in sequencing depth, sample composition, or the quality and quantity of material used for Hi-C library preparation, particularly given that the Hi-C data for these assemblies were generated from a mixed-infection sample. Taken together, these patterns suggest that *P. ovale wallikeri* retains a tightly clustered centromere organization, whereas *P. malariae* and *P. falciparum* may display multiple centromere-associated foci or weaker inter-centromeric interactions under the conditions sampled. While chromosomal interactions are not evident in PfML47, we see clear indications of interactions in the subtelomeric regions, highlighted with maroon boxes in **Fig. 1c**

Across *Plasmodium* species, conserved features of genome architecture include clustering of centromeres, telomeres, and virulence genes ^40–42^. However, the degree and functional implications of this clustering vary among species. In PowML03 and PmML03 especially, the local domain structure within each chromosome appears to differ markedly (**Fig 1a,b**). In addition to the clustering of centromeres, we observe distinct domains in the subtelomeric regions, usually associated with multigene families (discussed in detail below), indicating that spatial genome organization in these species might constrain the distribution of the multigene families. In *P. falciparum*, virulence gene clusters on different chromosomes colocalize in 3D influencing overall genome organization and correlate with their coordinated silencing ^41^. Beyond *Plasmodium*, *Babesia microti* genome shows a Rabl organization with subtelomeric virulence gene colocalization, whereas *Toxoplasma gondii* lacks virulence gene clustering and is instead dominated by centromeric associations ^42^.

The delineation of distinct chromosomal domains was further supported by analysis of GC content in PowML03 and PmML03, displaying an isochoric structure, with a marked transition between the relatively GC-rich internal chromosomal regions and the AT-rich subtelomeric regions. (**Fig. 1a,b**). This is also observed in *P. vivax*, *P. cynomolgi*, and *P. brasilianum* (**Extended Data Fig. 1f**). The pattern of GC content in PowML03 is similar to *P. vivax* and *P. cynomolgi*, while *P. malariae* is expectedly similar to *P. brasilianum,* which is considered to be nearly identical to *P. malariae* (**Extended Data Fig. 1f**) ^43^. *P. falciparum* shows an inverted pattern, where the extreme ends of the chromosome were GC rich compared to the more AT-rich core (**Extended Data Fig. 1f**). Furthermore, the centromeres of PowML03 and PmML03, which were identified from Hi-C contact maps, coincide with the steep localised drop in GC content observed within the core genome (**Fig. 1a,b, Supplementary Table 2**). Because of the low or indistinct cross-shaped interaction pattern in the Hi-C contact maps of *P. falciparum* assemblies, this characteristic drop in GC content, genomic intervals where GC content dropped below 5% across two consecutive 1000 bp bins, was used instead to infer the centromeric regions in *P. falciparum* (**Supplementary Table 2)**. This criterion when applied to Pf3D7 is concordant with centromere positions reported earlier ^38,39^ (**Supplementary Table 2)**. Other than the low GC content, the centromeric regions of PowML03 and PmML03 showed no discernible repeat patterns or intra- or inter-chromosomal sequence similarity, mirroring what is seen in *P. falciparum* ^38,44^.

When interpreting the described Hi-C contact maps, we should account for stage heterogeneity and the specimens used here for scaffolding the resultant assemblies. While PowML03 and PmML03 were each scaffolded with their respective Hi-C data, PfML47 was scaffolded using Hi-C data from another participant, MSC29 (**Extended Data Fig. 1a**). In such a case, while large scale interaction patterns are expected to be robust, finer sample-specific structural variations, especially in the subtelomeric regions, should be interpreted with caution. Similarly, stage composition can influence Hi-C signal with each stage presenting a different chromosomal configuration ^41,42^. The Hi-C library used to scaffold the *P. falciparum* assemblies was prepared from the enriched fraction of sample MSC29, which contains predominantly early asexual (ring-stage) parasites. On the other hand, *P. malariae* and *P. ovale wallikeri* Hi-C libraries were generated from enriched fractions that contained mixed erythrocytic stages, but largely represented by trophozoites. In *P. falciparum*, chromatin structure has been shown to change dramatically through the erythrocytic cycle, with the Hi-C contact maps of mid-stage trophozoites showing an open and intermingled chromatin conformation compared to early rings, and exhibiting increased inter-chromosomal interactions ^41^. Therefore, Hi-C data from a ring-stage enriched sample might underrepresent the configuration we see in trophozoites. In PowML03 and PmML03, Hi-C from a heterogeneous parasite mixture might blur stage-specific contact patterns and show broader, less sharply defined domains. Nonetheless, we do see clear interaction patterns in these two assemblies here, likely due to the high representation of trophozoites in the participants.

### Comprehensive Gene Annotation of High-Quality Genome Assemblies

Annotation of the improved assemblies with the Companion tool increased the number of coding genes and pseudogenes across nuclear, mitochondrial, and apicoplast genomes compared to the existing best reference annotations ^45^ (**Table 2, Fig. 2a-c, Extended Data Fig. 3a-c, Supplementary Table 3**). Companion uses Liftoff ^46^ to transfer annotations from an existing closely related references (PocGH01, PmUG01, Pf3D7), which resulted in annotations of 5171, 5226 and 5098 genes in PowML03, PmML03, and PfML47, respectively with high sequence identity and alignment coverage compared to the reference in exon regions. A further 1863, 1498 and 680 genes, respectively were annotated by the BRAKER2 ^47^ implementation in Companion that uses AUGUSTUS ^48^ and GeneMark ^49^ to perform *ab initio* gene prediction by integrating the information of protein homology using the translated amino acid sequences to reference databases into the prediction. Non-coding gene prediction in the assemblies was performed with Aragorn and Infernal implementations in Companion, in addition to the Liftoff associated annotation transfer, yielding 130, 122, and 161 genes, respectively. Companion-predicted pseudogenes accounted for the rest of the 952, 550, and 174 annotations respectively for PowML03, PmML03 and PfML47 (**Table 2**). Of these, 537, 264, and 85 pseudogenes respectively were conserved across the human-infecting *Plasmodium* species and suggest that they most likely represent annotation artefacts arising from sequencing or assembly errors, and need manual curation to correct, pending validation from additional evidence such as orthogonal sequencing data and transcriptomic support. In *P. falciparum*, in addition to the well-characterized *var* gene family, a number of additional loci were annotated as “hypothetical proteins” beyond the *var* genes close to the telomeric ends (**Extended Data Fig. 3h**). These short genes lack any identifiable domains. Long non-coding RNAs derived from subtelomeric regions have been reported previously ^50,51^. Overall, by integrating homology-based transfer with *ab initio* prediction, the assemblies of PowML03, PmML03 and PfML47 captured a total number of 8116, 7396, and 6113 genes respectively, comprising both conserved and lineage-specific genes.

**Table 2.** Companion annotation statistics of PowML03, PmML03 and PfML47 in comparison to the existing reference annotations. Gene families’ counts in brackets indicate (coding genes, pseudogenes). “Core” and “non-core” regions of the chromosomes were defined by the boundaries of the outermost position on each chromosome end of a 1:1 ortholog conserved across all human-infecting *Plasmodium* species, with the “non-core” regions corresponding to the sub-telomeric regions. “Unplaced” contig refers to unassembled scaffolds with known chromosomes, while “unlocalised” refers to those with unknown chromosomes. The categories of gene families displayed in the table are described in Methods.

|  |  | *P. ovale wallikeri* |  |  | *P. malariae* |  | *P. falciparum* |  |
| --- | --- | --- | --- | --- | --- | --- | --- | --- |
|  |  | PowML03 | Pow222 | PowCR01 | PmML03 | PmUG01 | PfML47 | Pf3D7 |
| Companion Tools | Liftoff / RATT, MetaEuk, GenomeTools (for Pow222) | 5,144 mRNA, 27 rRNA | 4,826 mRNA | --- | 5,203 mRNA, 23 rRNA | --- | 5,051 mRNA, 47 rRNA | --- |
|  | AUGUSTUS | 1,822 mRNA | 2,280 mRNA | --- | 1,450 mRNA | --- | 676 mRNA | --- |
|  | GeneMark.hmm3 | 41 mRNA | --- | --- | 48 mRNA | --- | 4 mRNA | --- |
|  | Pseudo | 952 pseudogenes | 479 pseudogenes | --- | 550 pseudogenes | --- | 174 pseudogenes | --- |
|  | INFERNAL / Companion (for Pow222) | 15 rRNA, 7 ncRNA, 23 snoRNA, 6 snRNA | 11 rRNA, 7 ncRNA, 21 snoRNA, 5 snRNA | --- | 10 rRNA, 8 ncRNA, 22 snoRNA, 6 snRNA | --- | 18 rRNA, 31 ncRNA, 28 snoRNA, 6 snRNA | --- |
|  | Aragorn | 79 tRNA | 75 tRNA | --- | 76 tRNA | --- | 78 tRNA | --- |
| Gene type | mRNA | 7007 | 7182 | 6228 | 6701 | 5942 | 5731 | 5318 |
|  | pseudogene | 952 | 479 | 715 | 550 | 637 | 174 | 158 |
|  | noncoding | 157 | 119 | 104 | 145 | 136 | 208 | 244 |
|  | rRNA | 42 | 11 | 17 | 33 | 26 | 65 | 65 |
|  | snoRNA | 23 | 21 | 23 | 22 | 20 | 28 | 41 |
|  | snRNA | 6 | 5 | 6 | 6 | 4 | 6 | 5 |
|  | tRNA | 79 | 75 | 58 | 76 | 68 | 78 | 78 |
|  | ncRNA | 7 | 7 | 0 | 8 | 18 | 31 | 55 |
| Gene location | core | 4989 | 4989 | 4511 | 5033 | 4810 | 4923 | 4927 |
|  | non-core | 3009 | 874 | 191 | 2295 | 1139 | 1072 | 684 |
|  | unlocalised/unplaced contigs | 39 | 1763 | 2302 | 0 | 699 | 26 | 0 |
| Gene families | *fam-m/fam-l* | --- | --- | --- | 720 (639, 81) | 679 (483, 196) | --- | --- |
|  | *pir* | 2062 (1731, 331) | 1774 (1584, 190) | 1189 (1189, 0) | 221 (193, 28) | 257 (137, 120) | --- | --- |
|  | *var* | --- | --- | --- | --- | --- | 99 (75, 24) | 105 (65, 40) |
|  | *rifin/stevor* | --- | --- | --- | --- | --- | 205 (187, 18) | 184 (157, 27) |
|  | *sica/stp1/surfin* | 95 (77, 18) | 129 (117, 12) | 78 (78, 0) | 75 (67, 8) | 166 (109, 57) | 10 (10, 0) | 10 (7, 3) |
|  | *etramp* | 10 (10, 0) | 10 (10, 0) | 11 (11, 0) | 9 (9, 0) | 8 (8, 0) | 14 (14, 0) | 14 (14, 0) |
|  | *exp* | 101 (89, 12) | 128 (116, 12) | 105 (105, 0) | 417 (348, 69) | 523 (314, 209) | 102 (95, 7) | 113 (88, 25) |
|  | *phist/resa* | 62 (56, 6) | 62 (61, 1) | 32 (32, 0) | 35 (35, 0) | 30 (29, 1) | 74 (63, 11) | 77 (58, 19) |
|  | *rbp* | 16 (13, 3) | 16 (12, 4) | 7 (7, 0) | 6 (4, 2) | 5 (3, 2) | 7 (7, 0) | 7 (5, 2) |
|  | conserved | 1276 (1138, 138) | 1364 (1310, 54) | 1696 (1696, 0) | 1286 (1191, 95) | 1272 (1271, 1) | 1284 (1261, 23) | 1256 (1256, 0) |
|  | hypothetical | 826 (659, 167) | 495 (485, 10) | 467 (467, 0) | 858 (774, 84) | 54 (54, 0) | 493 (455, 38) | 6 (6, 0) |

Subtelomeric regions of *Plasmodium* genomes typically harbor rapidly evolving, lineage-specific gene families associated with immune evasion and host–parasite interactions. These multigene family proteins were annotated at the sub-telomeric regions in our new assemblies, whereas in previous genome assemblies, contigs containing these gene families were largely unlocalized to chromosomes. In *P. ovale wallikeri* (PowML03), these regions were enriched in the *Plasmodium* interspersed repeat (*pir*) genes, the surfin-related subtelomeric protein 1 (*STP1*) family, the *SICAvar* family, and the *phist* family. We identified 2,138 (1807 protein coding, 331 pseudogenes) *pir* genes that comprise about 25% of the genome’s total gene content (**Table 2, Fig. 2a**). In contrast, *P. malariae* (PmML03) subtelomeric regions contained *fam-l* and *fam-m* genes, as well as *pir, STP1*, *SICAvar*, and *phist* gene families consistent with previous reports from other species and earlier incomplete assemblies (**Fig. 2b, Table 2**). *fam-m* and *fam-l* genes, which are unique to *P. malariae*, comprise 720 of the 7403 genes in the PmML03 assembly and are presumed to be involved in host-parasite interactions (**Table 2**). Genes termed “*Plasmodium* exported protein, unknown function” are enriched in the non-core regions of both PowML03 and PmML03, and are highly expanded in PmML03 numbering more than 400. In *P. falciparum* (PfML47), subtelomeric regions were dominated by the *var* gene family, together with *rifins, stevor*, *STP1*, *SICAvar*, and *phist* gene families as expected (**Fig. 2c, Table 2**).

In both PowML03 and PmML03, the distinct domains in chromosome organization observed in Hi-C contact maps, as well as patterns in GC content, are further reflected in the distribution of multigene families and single-copy orthologs (SCOs) or conserved genes (**Fig. 1,2**). SCOs and conserved genes across *Plasmodium* species, in particular those labelled “conserved”, were present almost exclusively within the core genome (**Fig. 2a-c**) within central chromosomal blocks in Hi-C contact maps. Genes from multigene families such as *pir*, *fam-m/l, STP1*, *SICAvar*, and *phist* were clustered in the subtelomeric regions, aligning with the outer domains of the Hi-C contact maps, with the lengths of these outer domains corresponding closely to the subtelomeric regions harboring these gene families. In contrast to the *var* genes in *P. falciparum*, we did not observe internal localization within the core genome of any of these multigene family members in *P. ovale wallikeri* and *P. malariae*. This partitioning reflects a “two-speed genome” model, described in several plant pathogens such as aphids, phytophthora and fungi ^52–56^, where a conserved core encodes essential parasite biology, while subtelomeric ‘adaptive zones’ harbour rapidly evolving multigene families involved in host-parasite interactions. The gene distribution provides complementary evidence for distinguishing core from non-core regions in the assemblies, and we defined the boundaries of the core genome for PowML03 and PmML03 by the outermost position on each chromosome end of a 1:1 ortholog conserved across all human-infecting *Plasmodium* species (**Fig. 2a,b, Supplementary Table 2**).

In both PowML03 and PmML03, the skewed AT content within the sub-telomeric regions across the chromosomes appears to be mainly driven by the intronic regions of the multigene families, as well as the intergenic regions (**Extended Data Fig. 3d-e**). In *P. falciparum*, the *var*, *stevor* and *rifin* mulitigene families contribute to a higher GC content compared to the rest of genes, especially within their exonic regions (**Extended Data Fig. 3f**). In addition to the gene body, intergenic regions of PowML03 and PmML03 were equally low in their GC content in the sub-telomeric regions compared to the core genome (**Extended Data Fig. 3g**).

### Multigene families

*P. ovale* spp. and *P. malariae* species exhibit striking expansions in gene families potentially linked to immune evasion and host interactions, particularly the *Plasmodium* interspersed repeat (*pir*) superfamily in *P. ovale* spp. and the *fam* genes in *P. malariae* ^1,2,26,28^. These expansions highlight their distinct evolutionary trajectories and studying them can provide insights into the mechanisms underlying chronic infection, immune evasion, and host adaptations in these neglected malaria species.

### Pirs in *P. ovale wallikeri* and *P. malariae*

The *Plasmodium* interspersed repeat (*pir)* gene family has undergone extensive lineage-specific expansions in *Plasmodium* species, giving rise to a large and heterogeneous repertoire, ranging from ∼200 in *P. chabaudi* ^57^, 170-224 *pirs* in *P. berghei,* ^58^ to over 2,100 in *P. ovale wallikeri* (PowML03) as shown here (**Table 2**). These expansions in PowML03, concentrated in subtelomeric regions, contribute to a ∼30% increase in genome size compared to other *Plasmodium* species (**Table 2, Fig. 2a**). *P. ovale wallikeri pir* genes were distributed across all chromosomes, showing no chromosome-specific clustering (**Fig. 2a**). Transcript lengths of predicted protein coding *pirs* range from 198 to 2664 bp, with exon counts from 1 to 7 (**Extended Data Fig. 4a**). Typically, *pir* genes lack a signal peptide, the N-terminal amino acid sequence that directs proteins to the secretory pathway (**Extended Data Fig. 4a**). They contain three exons: a short first exon, a longer second exon with a transmembrane (TM) domain, and a shorter third exon (**Fig. 3a,b, Extended Data Fig. 4a**).

**Fig. 3:**
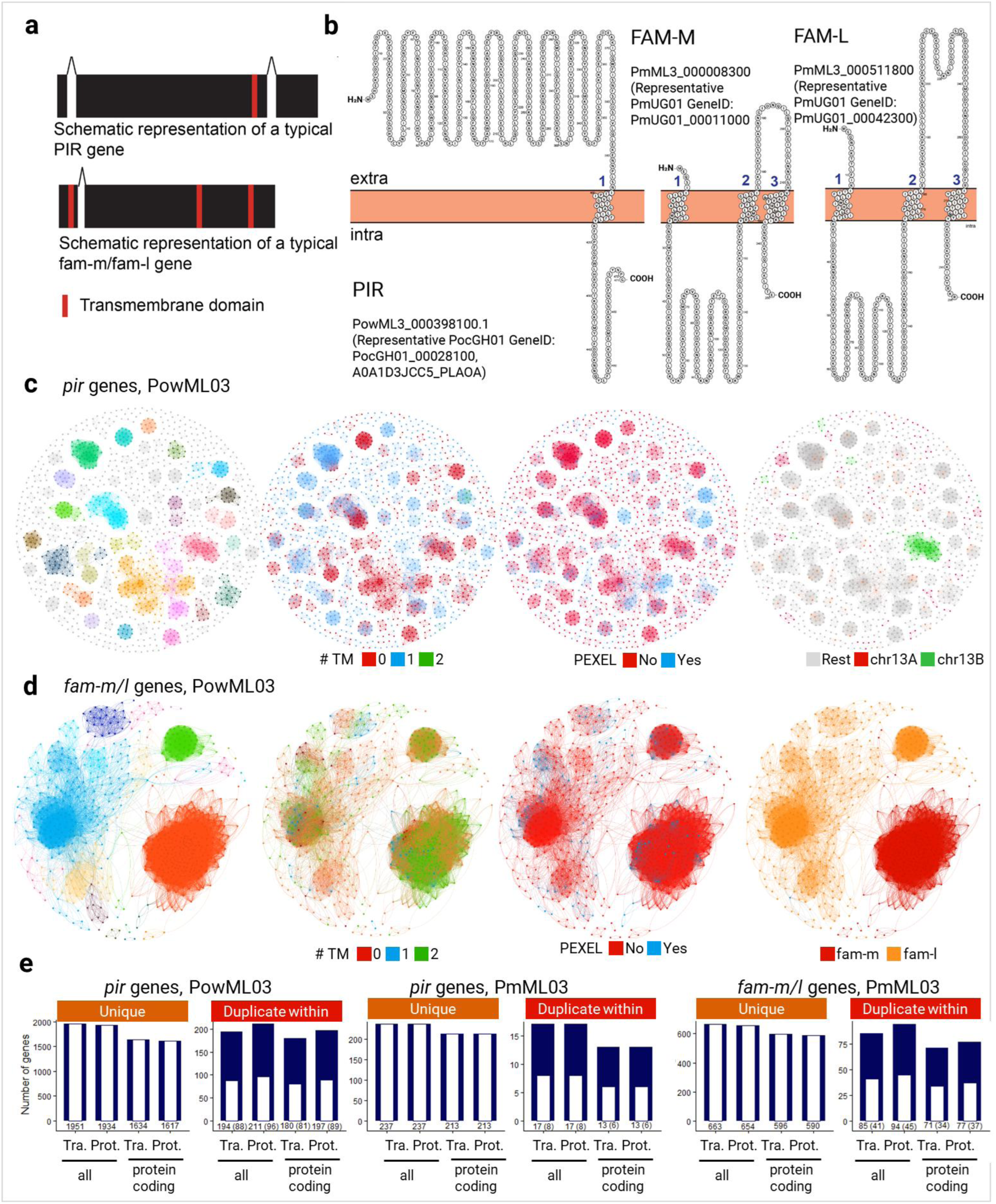
a, Typical exonic structures of *pir* genes in *P. ovale wallikeri* (PowML03) and *P. malariae* (PmML03), and *fam-m/fam-l* genes in *P. malariae* (PmML03) (black rectangles as exons, with red block indicating transmembrane domain). **b,** Example of topology of a typical PIR generated from Protter using the protein sequence ID corresponding to the sequence in *P. ovale curtisi* protein, PocGH01_00028100. **c,** Protein similarity network of PIRs in PowML03. Colored by module, by number of transmembrane domains (TM), by presence of relaxed PEXEL motif (RxLxx[EQD] or RxLx[EQD]), and by *pir* genes location in chr13 (indicated in **Extended Data** Fig. 5). **d,** Protein similarity network of the FAM-M/L proteins in PmML03. Colored by module, by number of transmembrane domains (TM), by presence of PEXEL motif (relaxed motif R.L..[EQD]), and by gene description. **e,** Plot showing the total number of genes grouped by whether transcript (Tra) or amino acid (Prot) sequence is unique or duplicated within the sample. White bars within quantify the number of unique sequences amongst the duplicated sequences. Data is shown for both “all” genes and only “protein coding” genes of the indicated families.

Protein similarity networks were constructed from all-versus-all BLASTP searches of *P. ovale wallikeri* (PowML03) amino acid sequences from all 1807 protein-coding *pirs*, retaining 1381 PIRS with ≥50 % identity over ≥100 aligned residues. The resulting networks were visualized in Gephi ^59^ using a force-directed layout to reveal relationships among these multigene family members (**Fig. 3c-f**). The *pir* family formed a diffuse network, with several distinct clusters, suggesting functional specialisation within the family. Furthermore, self alignment of the PowML03 subtelomeric regions revealed extensive sequence similarity across the non-core regions likely suggesting ongoing segmental duplication (**Extended Data Fig. 5a,d**). In addition, two distinct regions were evident on the uncharacteristically long *pir*-filled arm of chromosome 13, which appear to be distinct and dissimilar to the rest of the regions (**Extended Data Fig. 5d**). This is further reflected in the protein similarity network, where the subset of PIRs from the long arm of chromosome 13, grouped into a clear cluster, probably indicating local expansion events.

*P. ovale* and *P. malariae* encode all the components of the PTEX (*Plasmodium* translocon of exported proteins) machinery described in *P. falciparum* and *P. berghei*, which is involved in export of proteins to the host cell. A subset of PIR proteins contain the canonical PEXEL (*Plasmodium* export element) motif that is required for protein export beyond the parasitophorous vacuole membrane (**Extended Data Fig. 4a**), and are likely exposed on the surface of infected red blood cells, where they could play key roles in antigenic variation and immune evasion ^60,61^. The protein topology prediction suggests that the larger region before the TM-domain is presented on the cell surface while the shorter C-terminal portion of the protein is oriented toward the interior of the host cell, potentially important for trafficking to the RBC surface or for downstream signaling events within the infected red blood cell (**Fig. 3b**). Most *pir* genes contain a ∼164 bp exon located downstream of the transmembrane region, and the C-terminal region of PIR proteins shows increased sequence conservation, consistent with a conserved functional role. However, most PIRs lacked the PEXEL motif, suggesting that not all are exported, or involve non-PEXEL mediated export.

The pirs in *P. malariae*, also display an expanded repertoire, albeit much less than *P. ovale* or *P. vivax* (**Table 2, Fig. 2b, Extended Data Fig. 4b,c**), with 254 *pir* genes in PmML03 (226 protein coding, 28 pseudogenes). They display an exonic structure similar to those in *P. ovale wallikeri* and distributed chromosomal location (**Fig. 2b**, **Extended Data Fig. 4b**). Protein similarity network comprising *pir* genes from both PmML03 and PowML03 show distinct, non-overlapping clusters for each species, indicating substantial sequence divergence among pir genes across these two species (**Extended Data Fig. 4d**).

We assessed sequence redundancy within the *pir* gene family in PowML03 across the chromosomes by classifying the transcript and protein sequences into unique singletons and those duplicated within the specimen. We noticed sequence redundancy within PowML03 across the chromosomes, identifying 194/2145 *pir* gene transcripts, comprising 88 unique sequences that were present in multiple (2-4) identical copies, indicative of recent tandem or segmental duplication events (**Fig. 3e**). Likewise, within the *pirs* in PmML03, 17/254 (comprising 8 unique copies) occur in multiple (2-3) identical copies.

### fam-m, fam-l genes in *P. malariae*

The 720 fam-m/l genes were similarly concentrated in subtelomeric regions, along with other surface family genes, and show no chromosome-specific localisation (**Table 2, Fig. 2b**). Transcript lengths of protein coding fams range from 252 to 805 bp for fam-m and 95 to 937 bp for fam-l genes (**Fig. 3a, Extended Data Fig. 4e**). Both fam-m and fam-l genes range from one to four exons, with the most typical exonic structure (∼ 93% of the genes) being two exons with a short first exon containing a transmembrane domain and a long second exon with two TM domains. Only a subset of *fam-m* and *fam-l* proteins (28 and 43, respectively) are predicted to contain an N-terminal signal peptide (**Extended Data Fig. 4e**). Around 10% of the *fam-m/l* genes harbor the PEXEL motif and are likely exported to the host cell (**Extended Data Fig. 4e**) ^26^. The protein topology suggests a large fragment of the protein (between the first and second TM) is intracellular while a short segment between TM2 and 3 is presented on the cell surface with a short C-terminal tail within the cytoplasm (**Fig. 3b**). fam-m/l proteins show structural homology with PfRh5, which is key for RBC invasion, and they may have a role in host cell binding ^26^. Protein similarity networks were clustered from all-versus-all BLASTP searches of *P. malariae* (PmML03) protein-coding fam-m/fam-l genes (667), retaining sequence pairs with ≥50 % identity over ≥100 aligned residues. The network comprising 268 genes showed clear and well-separated clusters of fam-m and fam-l sequences, as expected based on family-level classification (**Fig. 3d**). Within the fam-l group, two major clusters were evident, suggesting the presence of distinct fam-m subfamilies or divergent lineages (**Fig. 3d**). No clustering patterns were associated with chromosomal location or number of transmembrane domains, indicating that *fam* gene similarity primarily reflects sequence divergence rather than genomic context or structural features (**Fig. 3d**). Similar to *pir* genes, within-genome redundancy of *fam* transcripts and derived amino acid sequences was notable. We identified 85/748 *fam-m/l* transcripts, comprising 41 unique sequences that were present in multiple (2-4) identical copies.

### Syntenic Analysis & Recombination within Sub-telomeric Regions

High quality assemblies from multiple specimens enabled us to assess synteny within and across *Plasmodium* species, using Circos and SyRI/plotsr ^36,62,63^(**Fig. 4, Extended Data Fig. 6–9**). The new reference assemblies, PowML03 and PmML03, were each scaffolded using Hi-C data derived from their respective specimens. The additional assemblies (PowML04, 08, 09 and PmML04, 07), however, were scaffolded using Hi-C data from the same specimens used for PowML03 and PmML03, respectively. In contrast, the *P. falciparum* assemblies, PfML46 and PfML47, were scaffolded using Hi-C data from a different participant, MSC29, which also yielded the PfML72 assembly (**Extended Data Fig. 1a**). Caution should therefore be exercised when interpreting several assemblies, as the Hi-C data originating from a different individual and developmental stage may influence contact patterns and thereby the final assemblies. Although minor scaffolding inaccuracies cannot be fully excluded, the overall correctness of the assemblies is supported by the strong conservation of synteny across core genomic regions, including those spanning scaffold joins (**Fig. 4a, Extended Data Fig. 6c-e, Extended Data Fig. 7a-c**). Contigs from earlier published references and other specimens from this study that could not be localised to chromosomes primarily map to the sub-telomeric regions of PowML03 and PmML03, and are typically AT-rich (**Extended Data Fig. 6d-f**). Two large unlocalised contigs in PowML04 and PowML09 correspond to the uncharacteristically long sub-telomeric region of chr13 in PowML03 (**Extended Data Fig. 6d)**.

**Fig. 4.**
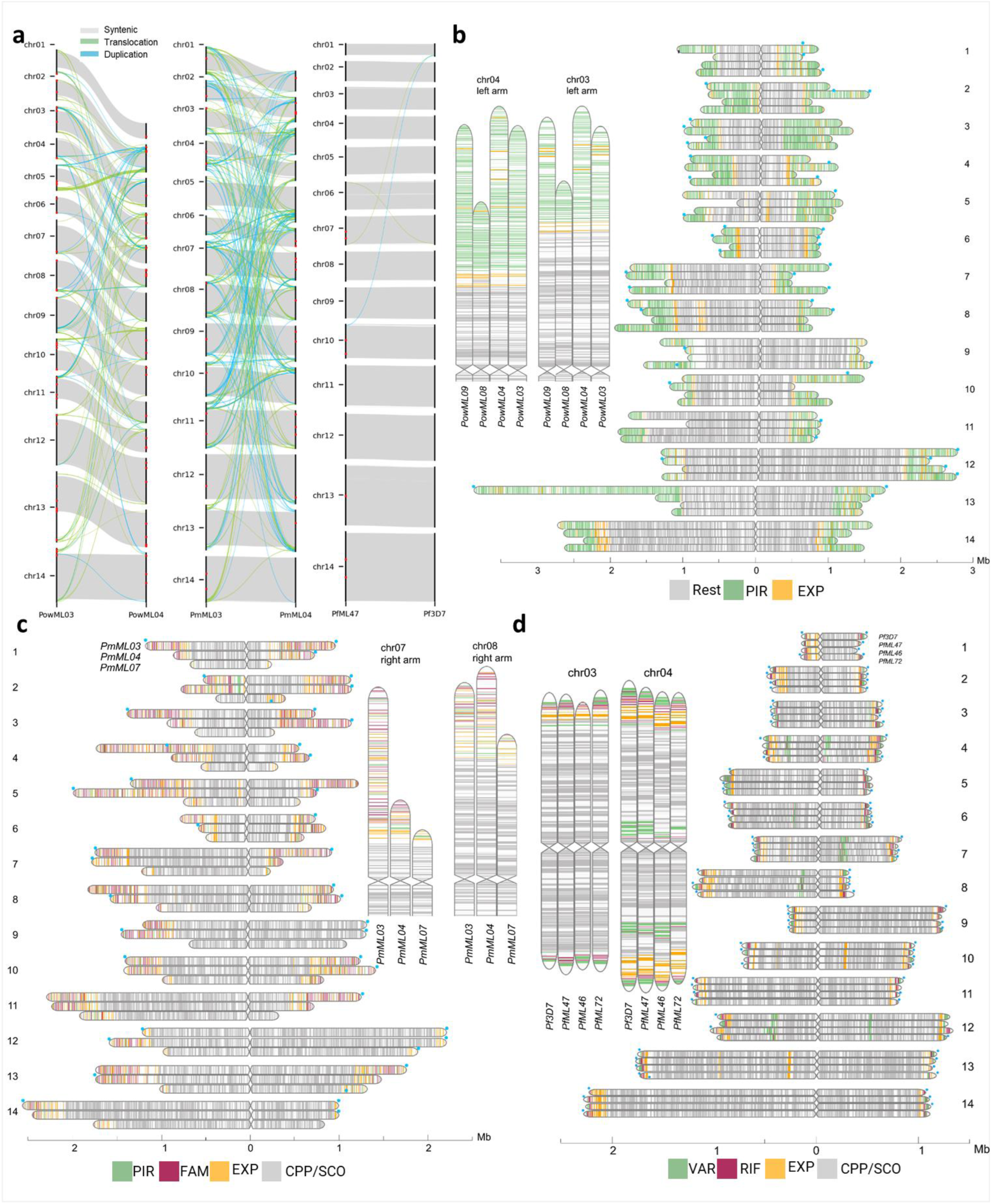
a, Synteny plots between the two best genome assemblies available for each species shows synteny within the core genome and a high level of sequence movement within the subtelomeric regions in *P. ovale wallikeri* and *P. malariae*. **b-d,** Comparison of gene annotations across all assemblies generated here show a high degree of gene order conservation within the core genome, which is lost towards the subtelomeric regions. A close up of chromosomal ends is shown in the insets. This syntenic order within the core genome is less apparent in PmML07 due to the comparatively poorer quality of the assembly. Synteny is also disrupted around the internal *var* genes found in chromosomes 4, 7, 8, and 12 of *P. falciparum*. Blue circles correspond to the presence of *Plasmodium* telomeric repeats, indicating likely chromosomal ends. Genes are grouped into the displayed gene families into categories as described in Methods.

In addition to the previously described PowML03, PmML03 and PfML47 annotations, the remaining assemblies of *P. ovale wallikeri*, *P. malariae* and *P. falciparum* were independently annotated by Companion ^45^, using the curated reference annotations from PocGH01, PmUG01, and Pf3D7, respectively to assess gene content **(Supplementary Table 3)**. In parallel, gene annotations from the best assembled genomes were transferred to these specimens, namely from PowML03 to PowML04,08,09, from PmML03 to PmML04, 07 and from Pf3D7 to PfML46,47,72 (**Extended Data Fig. 6a,b, Supplementary Table 4)** to enable cross-specimen comparisons using transferred gene IDs.

To assess sequence redundancy and conservation across isolates (i.e. the different genomes) of the expanded *pir* and *fam-m/l* multigene families, we classified genes across the annotations into three categories based on transcript and protein sequence identity: singletons, duplicates within a single isolate, and sequences shared across multiple isolates. Across PowML datasets, most protein and transcript sequences were identical across two or more isolates, followed by unique singletons. A pronounced fraction of transcript sequences showed redundancy both within, as described earlier for PowML03, and across the isolates (**Extended Data Fig. 6g-i**). For example, in PowML03, 1520 pir transcript sequences were shared across two or more isolates, 199 occurred as copies both within an isolate and in at least one other isolate, and 12 were present only as within-isolate copies (**Extended Data Fig. 6g**). The relative contribution of these categories varied across datasets, reflecting differences in assembly completeness annotation quality, with the best assemblies of *P. ovale wallieri* (PowML03 and PowML09) showing high proportion of shared sequences due to increased gene annotation. Remarkably, the proportion of sequences shared across isolates was higher in *P. ovale wallikeri* compared with *P. malariae*, indicating low sequence variation (**Extended Data Fig. 6g-i**). Recent population genomic studies indicate that both *P. ovale wallikeri and P. malariae* exhibit lower nucleotide diversity compared to *P. falciparum* and *P. ovale curtise* ^1,2^. However, a head-to-head comparison between these species, particularly from this geographic region, is lacking.

Extensive nucleotide sequence redundancy observed within the *pir* and *fam-m/l* multigene families complicates the interpretation of alignment-based synteny analyses like SyRI. As many of these genes are highly similar to each other or even occur as multiple identical copies dispersed across the genome, synteny tools struggle to resolve duplicated or interchangeable loci. To increase confidence in the accurate identification of multicopy sequences, we restricted interpretation to SyRI-defined blocks that are >10 kb in length (longer than most genes) and show >98% identity to each other, and only these high-confidence sequences were explored and visualised. We observe clear synteny across the core genome and extensive translocation and duplication in the non-core genome for both *P. ovale wallikeri* and *P. malariae* (**Fig. 4a, Extended Data Fig. 7a**).

Additionally, we used gene annotations to assess and quantify the dynamics between genomes in the AT rich sub-telomeric regions. To minimise artefacts arising from annotation transfer, stringent filtering criteria were used: only genes present as single copies (no two genes with identical nucleotide sequence of the gene body) in the assemblies PowML03 and PmML03 were considered, and only those successfully transferred by Liftoff with high confidence (coverage > 0.9 and sequence identity > 0.8) were retained for downstream analyses. We define a block as a contiguous set of orthologous genes (minimum of two genes) that maintain relative order in the chromosome of one genome but were relocated in another. We identified several blocks that have shifted positions between chromosomes, representing transposition events (**Extended Data Fig. 7d-g, Extended Data Fig. 8,9**). For example, in *P. malariae*, one block on chromosome 4 in PmML03 (spanning genes PmML03_000122500 - PmML03_000123400) is instead found on chromosome 11 in PmML04 (**Extended Data Fig. 9a**). This replicated block consists of 10 genes, with clear one-to-one correspondences between the source and target chromosomes, with synteny lost beyond this block. We find transposed blocks containing up to 30 genes in *P. ovale wallikeri* and 20 in *P. malariae* which typically include members of *pirs* and *fams*, suggesting potential functional implications for parasite adaptation and antigenic variation (**Extended Data Fig. 7d,e**). In *P. falciparum*, we find a maximum of 3 genes in these blocks, which consist mostly of rifins and exported proteins (**Extended Data Fig. 7f)**. Several more detailed examples for each species of block transposition are supported using both JBrowse ^64^ and SyRI ^36^ (**Extended Fig. 8,9**).

For a broader visualisation of the loss of synteny in the sub-telomeric regions in these species, Companion-annotated genomes were oriented by their predicted centromere location (**Fig. 4b-d, Supplementary Table 2**). For all within species comparisons, we see high level of synteny within the core genome, supported by a similar pattern of conserved genes (single copy orthologs across human infecting *Plasmodium* species, and genes labelled as “conserved *Plasmodium* protein” depicted with grey lines) (**Fig. 4b-d**). However, this synteny is lost outside the core genome. Similar gene rearrangements in the subtelomeric regions were described for *P. chabaudi* ^57^. Dynamic subtelomeric regions in *P. falciparum* are well–documented, especially in relation to the *var* gene family, which undergoes frequent ectopic recombination, facilitating antigenic variation ^65–67^. Comparable dynamics are evident here in *P. ovale wallikeri* and *P. malariae*, although the extent of this variation is striking given that these parasites were collected from the same village in a single malaria season. This observation underscores the capacity for rapid diversification in sub-telomeric regions even among closely related parasite populations in these two species.

Moreover, the extent of gene annotation discrepancies, upon transferring annotations from PowML03 and PmML03 to the other assemblies, within the multigene families compared to the conserved genes further points towards the extensive recombination events within the sub-telomeric regions, highlighting the rapidly evolving multigene families (**Extended Data Fig. 6a,b**).

### Other gene families, drug resistance markers, exported proteins

In addition to the two major gene families analysed above, we also surveyed other multi-copy and antigenic gene families across our assemblies, including SURFIN/STP1, MSPs, and genes commonly used as markers of invasion or sexual-stage biology (e.g., Pfg25/27). The assemblies PmML03 and PowML03 exhibit a dramatically expanded repertoire of SURFIN/STP1-type genes (∼90-120 copies), in sharp contrast to the ∼10-12 copies typical of *P. falciparum* (**Table 2**). SURFIN/STP1 proteins are implicated in host–parasite interactions, cytoadherence, and immune evasion. Similarly, substantial copy-number variation in MSPs and Pfg25/27 genes was observed in both *P. malariae* and *P. ovale wallikeri*. For example, we detect 14 tandem copies of MSP3 in PowML03 (chromosome 10) and 13 tandem copies in PmML03 (chromosome 6) (**Extended Data Fig. 10a)**. MSP3 family members play roles in merozoite invasion and are known vaccine targets. The differences are equally striking for Pfg25/27, a key marker of gametocytes crucial for gametocytogenesis (**Extended Data Fig. 10b)**. While *P. falciparum* (Pf3D7) and *P. ovale wallikeri* (PowML03) encode only 1–2 copies, PmML03 contains a large expansion of 25 copies (22 in PmML04 and PmUG01, 15 in PmML07) all arranged consecutively on chromosome 14 (**Extended Data Fig. 10b-c)** in addition to the single copy on chromosome 4 that is more closely related to the *P. falciparum* and *P. ovale* spp. orthologs. Such expansions suggest that *P. malariae* and *P. ovale wallikeri* may depend on a far broader set of exported or surface-exposed antigens. This could reflect differences in their strategies for immune modulation, chronicity, or host cell tropism. These patterns also highlight important implications for future diagnostics, where gene family expansion could influence molecular assay performance.

We also assessed known drug-resistance associated loci in *P. falciparum*. The triple *dhfr* mutant haplotype N51I/C59R/S108N, widely prevalent in Mali and often found close to 100%, was present in all three *P. falciparum* specimens sequenced here (**Extended Data Fig. 10d)**. This is consistent with regional patterns of sustained SP drug pressure for intermittent preventive treatment in pregnancy. We did not observe evidence of amplifications, deletions or copy number variations at mdr1 or crt loci or plasmepsins. Two of the three *P. falciparum* specimens (PfML47 and PfML72) harbor the CVIET (chloroquine resistant) haplotype across residues 72-76, along with additional mutations A220S, Q271E, I356T, and R371I, which have been shown to contribute to the resistance phenotype ^68,69^ (**Extended Data Fig. 10e)**. PfML46 and PfML72 carry the K189T mutation in *kelch13*, a common non-propeller polymorphism identified across Africa and Asia ^70^. This mutation has been shown to be associated with a longer parasite clearance time in patients treated with Artemisinin-based Combination Therapies (ACTs) ^71^. On the other hand, PfML47 isolate carries the S436A and G437A mutations in *dhps*, which are implicated in conferring resistance to sulfadoxine, particularly when occurring alongside other *dhps* or *dhfr* mutations ^72^. While none of the *P. ovale wallikeri* or *P. malariae* specimens from this study harbored mutations in the above positions (except for the A437 residue in *dhps* existing as uniformly wild-type), data on drug-resistance markers in these non-*falciparum* species remain scarce. Genomic studies of *P. malariae* and *P. ovale* isolates across Africa and Asia identified mutations associated with reduced pyrimethamine susceptibility ^23,73^. There is growing concern over emerging resistance, with a recent study from Mali identifying unusually high IC50 values for chloroquine, lumefantrine and artemether against *P. malariae* isolates, which are the current frontline drugs against *P. malariae* infections ^74^. These reports highlight the need for comprehensive surveillance and resistance studies of these neglected species.

## Discussion

In this study, we generated substantially improved reference assemblies for *Plasmodium ovale wallikeri* (PowML03), *Plasmodium malariae* (PmML03) using PacBio ULI HiFi long read sequencing and Hi-C from very little starting material from natural infections. Producing reference-grade genomes directly from low-input clinical material bypasses the need for lab culturing that can be associated with artefacts and adaptations ^75^. These assemblies offer more complete annotations of coding genes, pseudogenes, and non-coding elements across the nuclear, mitochondrial, and apicoplast genomes.

The genome of *P. brasilianum*, a simian-infecting species considered genetically and biologically close to *P. malariae*, has been recently sequenced and assembled using PacBio from *ex vivo* schizont material ^43^. Despite the significant amount of starting material and similar long-read technique, the assembly is similar to the existing PmUG01 and incomplete within the sub-telomeric regions. This highlights the utility of Hi-C data to obtain complete assemblies that better capture the full genomic complexity of these species, especially because of the repetitive nature of the sub-telomeric region genes. Hi-C data enabled scaffolding of 50 and 23 hifiasm contigs from PowML03 and PmML03, respectively into the 14 chromosomes of the final assembly. Altogether, PowML03 and PmML03 constitute the most complete genomes to date for *P. ovale wallikeri* and *P. malariae*.

These resources also open the door to more refined population, comparative, and functional genomics. Population genomics of these species remains in its infancy. Recent work has found limited structure in *P. ovale* spp. populations but evidence for differentiation in *P. malariae* ^1,2^. The improved annotation sets and genomes will be especially valuable for designing accurate species-specific assays to better detect and monitor *P. ovale* and *P. malariae* in genomic surveillance programmes, an urgent need given their underappreciated epidemiological impact. Sexual development is poorly characterised for *P. ovale* spp. and *P. malariae*, especially with regard to regulators (e.g. AP2-G, GDV1), and developmental niches. These gaps underscore the need for stage-resolved single-cell studies to uncover conserved and species-specific regulatory strategies across the life cycles. The large, highly repetitive subtelomeric gene families we’ve identified here may complicate read mapping and quantification. Standard short-read RNA-seq methods often collapse or misassign expression among paralogues, masking true transcriptional diversity. Future projects may benefit from long-read transcriptomics, repeat-aware mappers, and curated gene family databases, to accurately resolve expression patterns across expanded multigene families. The chromosome-scale assemblies generated in this study will be a valuable foundation for such work, providing accurate gene models and complete subtelomeric regions for interpreting transcriptional dynamics in future bulk and single-cell transcriptomic analyses.

The improved resolution of long repetitive tracts here enabled our investigation of large structural variants and copy-number changes. Our findings reinforce a common theme across *Plasmodium* species, which display a highly conserved and syntenic core genome in contrast to highly dynamic subtelomeric regions enriched in rapidly evolving multigene families. We see pronounced lineage-specific expansions, with *pir* genes in *P. ovale* spp. and *fam-l/fam-m* genes in *P. malariae*, alongside *stp1*, *sicavar* and *phist* families, mirroring the compartmentalisation seen in *P. falciparum* and *P. knowlesi*, where subtelomeric diversification drives immune evasion while the core genome maintains essential functions. Sharp transitions in GC content and conserved orthologue boundaries further underscore this stable-core, flexible-subtelomere architecture. Although the mechanisms underlying the extensive subtelomeric dynamism in *P. malariae* and *P. ovale* spp. remain unclear, they echo and potentially exceed patterns in *P. falciparum*, where *var* gene diversification is driven by frequent ectopic recombination and sequence exchange during mitosis ^67,76^. This raises the possibility that processes that shape *var* dynamics may also shape *pir* and *fam-m/l* evolution. Given that mutually exclusive expression of *var* genes underpins antigenic variation in *P. falciparum*, the presence of large subtelomeric families in *P. malariae* and *P. ovale* spp. suggests that similar regulatory mechanisms may operate, highlighting *P. malariae* and *P. ovale* spp. as promising systems for future investigation into the mechanisms driving this sub-telomeric dynamism. Future transcriptomic and epigenomic studies, particularly at single-cell resolution, will be critical for determining whether these expanded repertoires are actively regulated, stage-specific, or partly pseudogenised. Their large numbers also raise intriguing evolutionary questions. Expansions in *P. ovale wallikeri* and *P. malariae* may reflect broader host ranges or cross-species transmission potential, as suggested by reports of *P. brasilianum* in primates and *P. ovale*-related parasites in chimpanzees. Dissecting the functional and regulatory diversity within these multigene families will be central to understanding antigenic variation, chronic infection, and host adaptation in these understudied parasites.

Together, these assemblies advance both the technical and biological frontiers. Technically, they demonstrate that ULI and long-read sequencing can generate high-quality *Plasmodium* references from minute DNA quantities, a crucial feature when working with scarce clinical samples. Biologically, they highlight conserved genome organisation principles across *Plasmodium* species while revealing unique lineage-specific adaptations. Moving forward, these resources will underpin comparative, functional and population level studies, enable improved molecular diagnostics, and strengthen collaborative initiatives such as the The Pan African Vivax and Ovale Network (PAVON).

## Methods

### Genomic material and assembly generation

#### Sample origin and processing

All sample collections and usage were done in accordance with the guidelines of the Research Ethics Committee and the Nagoya Protocol.

Study participants aged 6-19 with asymptomatic malaria or those admitted to clinics with uncomplicated malaria infections from Faladie, Mali were included in the study. For this analysis, participants with *P. ovale*, *P. malariae* or *P. falciparum*, either as a single or mixed infection with other species, as determined by light microscopy, were included in the study. Informed consent was taken from the patients to obtain 6 mL of blood. Blood was drawn into a CPDA vacutainer tube. Samples were processed according to the protocol described here ^77^. Briefly, 3 mL of blood from the CPDA tube was mixed with 5mL of SA buffer (10 mM Tris-HCL, 166 mM NaCl, 10 mM Glucose, pH 7.37) and passed through a chilled MACS LS column to obtain late asexual stages and gametocytes. The flowthrough was passed twice through pre-PBS-washed Plasmodipur filters to remove leukocytes. This flowthrough was further washed with PBS before 100 µl of the pellet was subjected to Streptolysin-O lysis to remove uninfected RBCs, to obtain the FT-SLO fraction. We observed variable lysis with SLO across the study participants, thus ending up anywhere from ∼5 µl to ∼100 µl of the fraction. The remaining flowthrough was lysed with 0.2% saponin, washed thrice with PBS before pelleting to obtain the FT-SAP fraction. The enriched fractions were profiled for single cell transcriptomics (not pertaining to this study) and the remaining material, if any, was stored at −80°C until DNA extraction. The fractions used for generating the assemblies are listed in **Extended Data Fig. 1a**.

#### DNA extraction and Sequencing

High molecular weight (HMW) DNA was extracted from enriched parasite fractions (**Extended Data Fig. 1a**) using Qiagen’s MagAttract HMW DNA extraction kit on a KingFisher™ Apex Automated Extraction System from Thermo Fisher Scientific, and eluted in a volume of 100 μl. HMW DNA was sheared using the Covaris g-TUBE by passing the DNA through the g-Tube twice at 3500 rpm to obtain ∼10 kb fragments required for library preparation. DNA quantity measured with a Qubit Fluorometer using Qubit’s dsDNA High Sensitivity Assay kit. Fragment size distribution was evaluated on the Agilent Femto Pulse system (**Extended Data Fig. 1a**). In case of presence of shorter DNA fragments, sheared DNA was further purified and concentrated using 3.7X Ampure XP SPRI to remove fragments below 3kb.

PacBio HiFi DNA sequencing libraries were constructed according to the manufacturers’ instructions. Both lowInput (LI) and Ultra LowInput (ULI) libraries were prepared using PacBio SMRTbell® Express Template Prep Kit 2.0 and PacBio SMRTbell® gDNA Sample Amplification (noting only ULI libraries were successful, as described below).

Of the 25 samples processed using the above DNA extraction and shearing, 7 samples (those with >250ng sheared product in a volume of 46 µl) underwent library preparation with the LI protocol and the other 18 samples (<250ng sheared product, 46 µl) underwent the ULI library protocol. We attribute this disparity in quantity to the varying amounts of host white blood cells and DNA across the sampled fractions that were retained in the MACS column or were not filtered out by the Plasmodipur columns. In the first batch, 7 LI samples and 11/18 ULI samples were multiplexed together on a single SEQUEL IIe flow cell each. The other 7 ULI samples were sequenced in a separate batch, sharing one ⅕ of a Revio flow cell. *Plasmodium* genomes were typically 20-30 Mb and these were multiplexed to generate much higher coverage than the 30x required for assembly in order to overcome substantial human DNA contamination. All 7 LI samples and 10/18 ULI samples failed due to high human contamination and thus low parasite content. However, the remaining 8 ULI samples produced sufficient parasite data, resulting in assemblies with high coverage.

Hi-C data were generated from SLO fractions of participants MSC23 (a *P. ovale wallikeri* infection) and MSC29 (mixed infection with *P. malariae* and *P. falciparum*). The SLO fractions for these two samples underwent minimal lysis upon incubation with the SLO enzyme, and ∼200 µl and 100 µl of the samples respectively were frozen and available for Hi-C. We used the Arima2 kit and sequenced these two samples on an Illumina NovaSeq 6000 instrument using 1/8th of a lane, yielding ∼22.74 million reads for MSC23 and ∼94.72 million reads for MSC29. To provide an estimate of the amount of material present, the volumes of samples used for Hi-C correspond to approximately ∼64 ng and ∼82 ng of HMW DNA extracted from comparable sample volumes.

#### Curation and assembly

Quality filtered and deduplicated PacBio HiFi reads were first mapped to the human reference genome (GRCh38.p14) using minimap2 (v2.27) with the option ‘-ax map-hifi’. Unmapped reads representing possible *Plasmodium* reads were extracted using samtools with the ‘-bf 0×4’ flag. These unmapped reads were assembled into contigs with hifiasm (Cheng et al., 2021). The resulting assemblies were scaffolded with Hi-C data from participants MSC23 and MSC29 using YaHS ^33^. MSC23 (*P. ovale wallikeri* infection) was used to scaffold PowML03, 04, 08 and 09, while MSC29 (mixed infection with *P. malariae* and *P. falciparum*) was used to scaffold the *P. malariae* and *P. falciparum* assemblies. Manual curation and checks for contamination were done as described previously ^78^. Mitochondrial and apicoplast genomes were assembled using MitoHiFi ^34^.

### Analysis and Visualization

#### Identifying *P. ovale* species

To check the species identity of the *P. ovale* spp. specimens, we examined single nucleotide differences from the *cox1* gene locus upon mapping the raw reads of the *P. ovale* spp. specimens onto the published *P. ovale curtisi* PocGH01, PlasmoDB v68 and *P. ovale wallikeri* Pow222 ^28^ reference genomes with minimap2 (v2.27) with the option ‘-ax map-hifi’. We confirmed that all *P. ovale* we sequenced were highly genetically similar to *P. ovale wallikeri*.

#### Identification of centromeric coordinates and core genomes

Centromeric coordinates were inferred based on local GC-content profiles. For *P. ovale wallikeri* and *P. malariae* assemblies, which contain a relatively rich GC core, centromeres were identified by a sharp reduction in GC content within this region. This distinct drop was not observed visually in *P. falciparum* due to its overall high AT-richness. For *P. falciparum* assemblies, the centromeric regions were therefore defined as genomic intervals where GC content dropped below 5% across two consecutive 1000bp bins. The boundaries of the core genome for *P. ovale wallikeri* and *P. malariae* assemblies were defined by the outermost position at each chromosomal end of a 1:1 ortholog conserved across all human-infecting *Plasmodium* species. As complementary evidence, gene content and GC content was examined across all these assemblies, and these align with the boundaries so identified.

#### Annotation with companion and annotation transfer

The Companion tool was used to annotate the complete assemblies, including chrA, chrM and any unlocalised contigs (**Supplementary Table 3**). We used the *P. ovale curtisi* PocGH01 annotation for PowML03, 04, 08, 09; the *P. malariae* PmUG01 annotation for PmML03, 04, 07; and the *P. falciparum* Pf3D7 annotation for PfML46, PfML47 and PfML72. Companion was used without using transcript evidence or pseudochrome contiguation, and using default settings (“No, do not use reference protein evidence”, “Yes, perform pseudogene detection”, “Yes, use BRAKER for structural annotation”, “No, use Liftoff to transfer reference gene models”). In addition to the Companion derived annotations, for comparison of transfer between specimens, Liftoff was used to transfer annotations from PowML03/PmML03/Pf3D7 to the other specimens with “-copies” option to look for extra gene copies in the target genome, and default settings for the rest of the options (**Supplementary Table 4**).

The categories of gene families tabulated in the figures and table comprise genes whose description terms include the following text: fam-m/fam-l - “fam-m protein” or “fam-l protein”; pir - “PIR protein” or “Plasmodium vivax Vir protein, putative”; var - “*PfEMP1*” var genes; rifin/stevor - “Subtelomeric Variable Open Reading frame, putative” or “Rifin, putative”; sica - “SICA C-terminal inner membrane domain containing protein, putative” or “SICA_C domain-containing protein”; stp1 - “STP1 protein”; surfin - “surface-associated interspersed protein” or “SURFIN” ; etramp - “early transcribed membrane protein” or “Malarial early transcribed membrane protein (ETRAMP), putative”; exp - “Plasmodium exported protein, unknown function” or “Plasmodium exported protein (hyp*), unknown function”; phist - “*PHIST*”; resa - “Plasmodium RESA N-terminal, putative” or “PRESAN domain-containing protein”; rbp - “reticulocyte binding protein*” genes; conserved - “conserved Plasmodium protein, unknown function” or “conserved protein, unknown function” or “conserved Plasmodium membrane protein, unknown function”; hypothetical - “hypothetical protein” or “hypothetical protein, conserved”. In Fig. 4 an Extended Data Fig. 3, the genes are grouped into the displayed gene families based on the description terms including the following text: FAM or fam-m/l - “fam-m protein” or “fam-l protein”; PIR - “PIR protein” or “Plasmodium vivax Vir protein, putative”; VAR - “*PfEMP1*” var genes; RIF or RIF/STEVOR - “Subtelomeric Variable Open Reading frame, putative” or “Rifin, putative”; EXP - the following exported proteins, “SICA C-terminal inner membrane domain containing protein, putative”, “SICA_C domain-containing protein”, “STP1 protein”, “early transcribed membrane protein”, “Plasmodium exported protein, unknown function”, “Plasmodium exported protein (hyp*), unknown function”; “*PHIST*”; “Plasmodium RESA N-terminal, putative”, “PRESAN domain-containing protein”, “reticulocyte binding protein*”; CPP - “conserved Plasmodium protein, unknown function” or “conserved protein, unknown function” or “conserved Plasmodium membrane protein, unknown function”; HYPO - “hypothetical protein”; HYPOC “hypothetical protein, conserved”.

#### Identification of dynamic synteny blocks

Companion annotations of PowML03 and PmML03, together with PlasmoDB v68 Pf3D7 annotations and Liftoff-derived annotations for the remaining assemblies (PowML04,08,09, PmML04,07, and PfML46,47,72), were used for this analysis (**Supplementary Table 3 and 4**). Gene coordinates (chromosome, start, end, gene ID) were extracted for the regions of interest. Syntenic relationships between regions were inferred by identifying genes that retained the same gene identifier between source and Liftoff-target assembly annotations (e.g., annotated by Companion in PowML03, and their corresponding transferred annotations in PowML04). Only single-copy genes (genes that don’t share 100% sequence identity with any other gene) in the source (PowML03, PmML03, Pf3D7), and those that transferred with ≥0.9 coverage and ≥0.8 sequence identity were retained. Blocks of genes were defined as contiguous sets of orthologs maintaining relative order in one chromosome of a sample but relocated in another sample. Synteny plots were generated using R (4.1.2) with ggplot2, focusing on the blocks of interest flanked by a few surrounding genes.

#### Construction of Gene Similarity Networks

To examine sequence-level relationships among multigene families, we constructed protein similarity networks using an all-versus-all BLASTP approach. The fasta file of protein sequences of interest was used to build a BLAST protein database, and an all-versus-all BLASTP search was performed with an e-value threshold of 1×10⁻⁵. Pairwise connections (edges) between proteins were retained if they exhibited ≥50 % amino-acid identity over ≥100 aligned residues. Self-hits were removed. The resulting edge and node tables were imported into Gephi for network visualization (**Fig. 3c,d, Extended Data Fig. 4c,d**). Layouts were computed using the Fruchterman–Reingold force-directed algorithm to highlight clusters of closely related proteins. Nodes were annotated with gene family identity, chromosomal location, and predicted number of transmembrane (TM) domains, allowing visual comparison of family-specific and structural clustering patterns.

#### Tools for visualization

Dotplot-based comparative analysis (**Extended Data Fig. 2a**) was done to visualise collinearity using the web application D-GENIES ^35^ by aligner minimap2 v2.28 and using the “Many repeats” option for repeatedness. For the dotplot-based comparative analysis of the non-core regions of the assemblies (**Extended Data Fig. 5**), paf (Pairwise mApping Format) files were generated using minimap2 with the options “-k15 -w10 -m40 --cs”. The resulting files were filtered to remove alignments with less than 10,000 bp and then visualised using D-GENIES ^35^. For gene based syntenic visualization (**Extended Data Fig. 7**), the Circlize package in R was used. The chromoMap package was used for chromosomal level visualization of GC density (**Extended Data Fig. 1f**), coverage (**Extended Data Fig. 2d**) and telomeric repeat density (**Extended Data Fig. 2i**), across the chromosomes. Syntenic regions and structural variations between the assemblies were identified using SyRI and visualised using plotsr package ^36,62^ (**Fig. 2a-c, Extended Data Fig. 1g, Extended Data Fig. 6a)**. Protein sequence structures of select genes were obtained through Protter (Fig. 3) ^79^. The remaining plots were generated in R using ggplot2. Packages and their version used to generate the data, tables and figures are listed below.

### Tools and versions

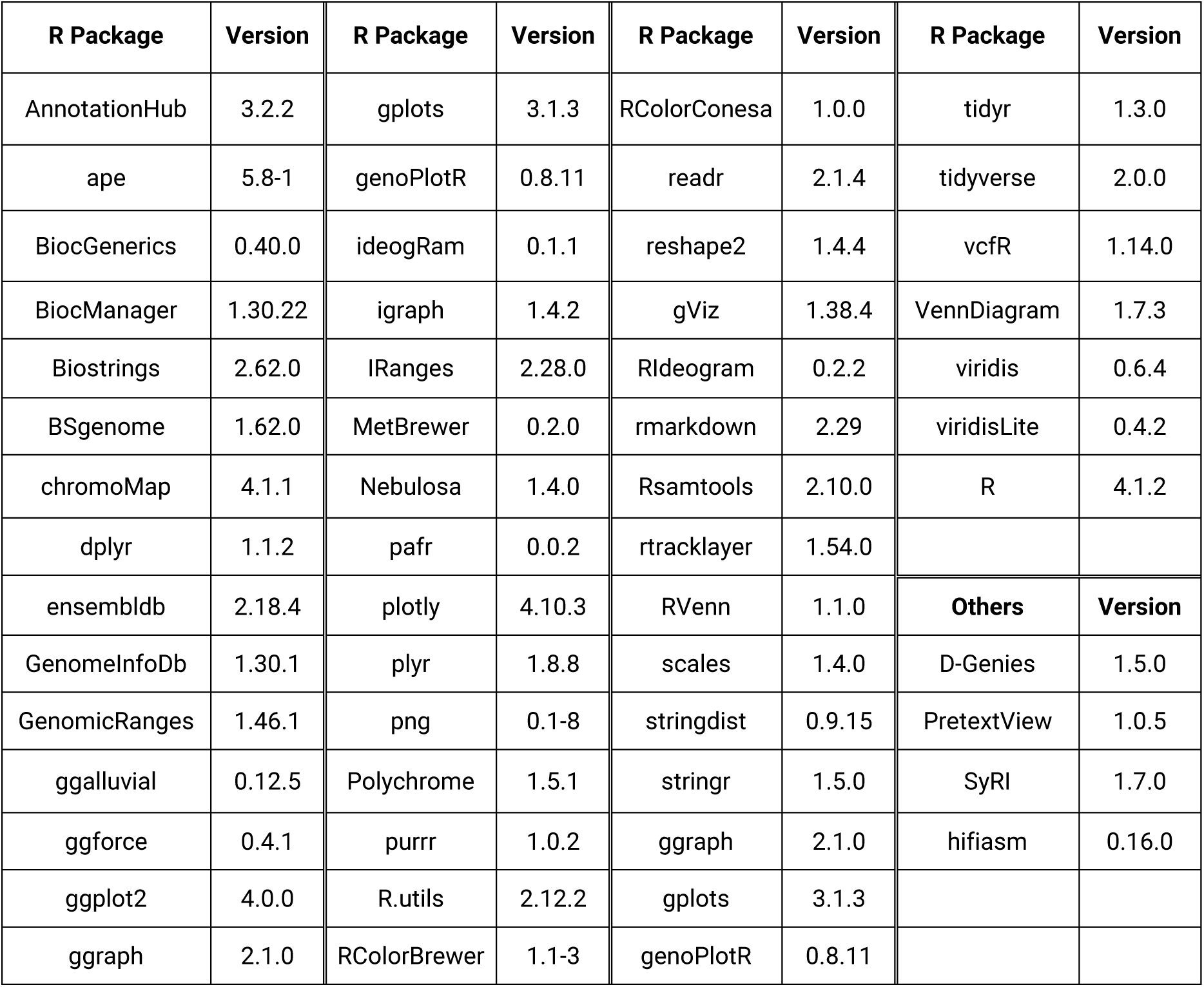

#### Data availability

The raw sequencing data is available at SRA under IDs ERS16235894-99, ERS18104290 and ERS18104302 and the assemblies are deposited in ENA with project accession IDs, PRJEB84978-86 (ERP168442-50), PRJEB82714 (ERP166389). Source data corresponding to the figures and tables are provided with the manuscript, and deposited, along with all the assemblies and annotations, at 10.5281/zenodo.18391093.

## Code availability

All the scripts and related data corresponding to figures and table generation are available at 10.5281/zenodo.18391093.

## Source data

Source data is provided in the supplementary files.

## Supplementary information

**Supplementary Table 1**

Excel document containing the mapping and assembly statistics

**Supplementary Table 2**

Excel document containing information pertaining to each assembly regarding number of gaps, telomere repeat regions, core genome and centromeric coordinates.

**Supplementary Table 3**

Excel document containing the gene summary tables from Companion derived annotations, using PocGH01 as reference for PowML03/04/08/09, PmUG01 as reference for PmML03/04/07 and Pf3D7 for PfML46/47/72.

**Supplementary Table 4**

Excel document containing the gene summary tables from Liftoff derived annotations (Annotations Liftoff from Pow222 to PowML03/04/08/09, from PowML03 to Pow222, PowML04/08/09, from PmUG01 to PmML03/04/07, and from PmML03 to PmML04/07.

## Acknowledgements

This work was supported by a Medical Research Council research grant (MR/S02445X/1) to MKNL, which supports SD. This work was also supported by Wellcome core funding to the Wellcome Sanger Institute (Grant 206194/Z/17/Z), which supports MKNL and JCR. Part of the field work in Mali was supported by the BMGF (INV-001927) through RCA No 20_356 between Sanger and USTTB. We thank the staff of the Wellcome Sanger Institute Scientific Operations for their contribution to library preparation and sequencing. We also thank the team that runs the clinic in Faladie, Mali for their contributions to sampling natural infections as well as the study population.

## Author information

None

## Contributions

MKNL, AMT and AAD conceptualized and supervised the study. AD, SS, DO and AAD supervised the clinical aspects of the study including recruitment of study participants. MKNL, AMT and AM provided guidance and supervision of key aspects of the study. SKD and JR were involved in sample processing from study participants. SKD and FT prepared HMW DNA for library preparation and sequencing. SKD analysed data, wrote the first draft, and edited subsequent drafts with input from all the co-authors. AM provided guidance on key analysis of the data. SKD, AM, JR, AMT and MKNL were involved in critical reading of the manuscript. SKD, DLP, YS, MUS, JT, TM and JW were involved in the curation of the assemblies.

## Extended Data

**Supplementary Table S1:**
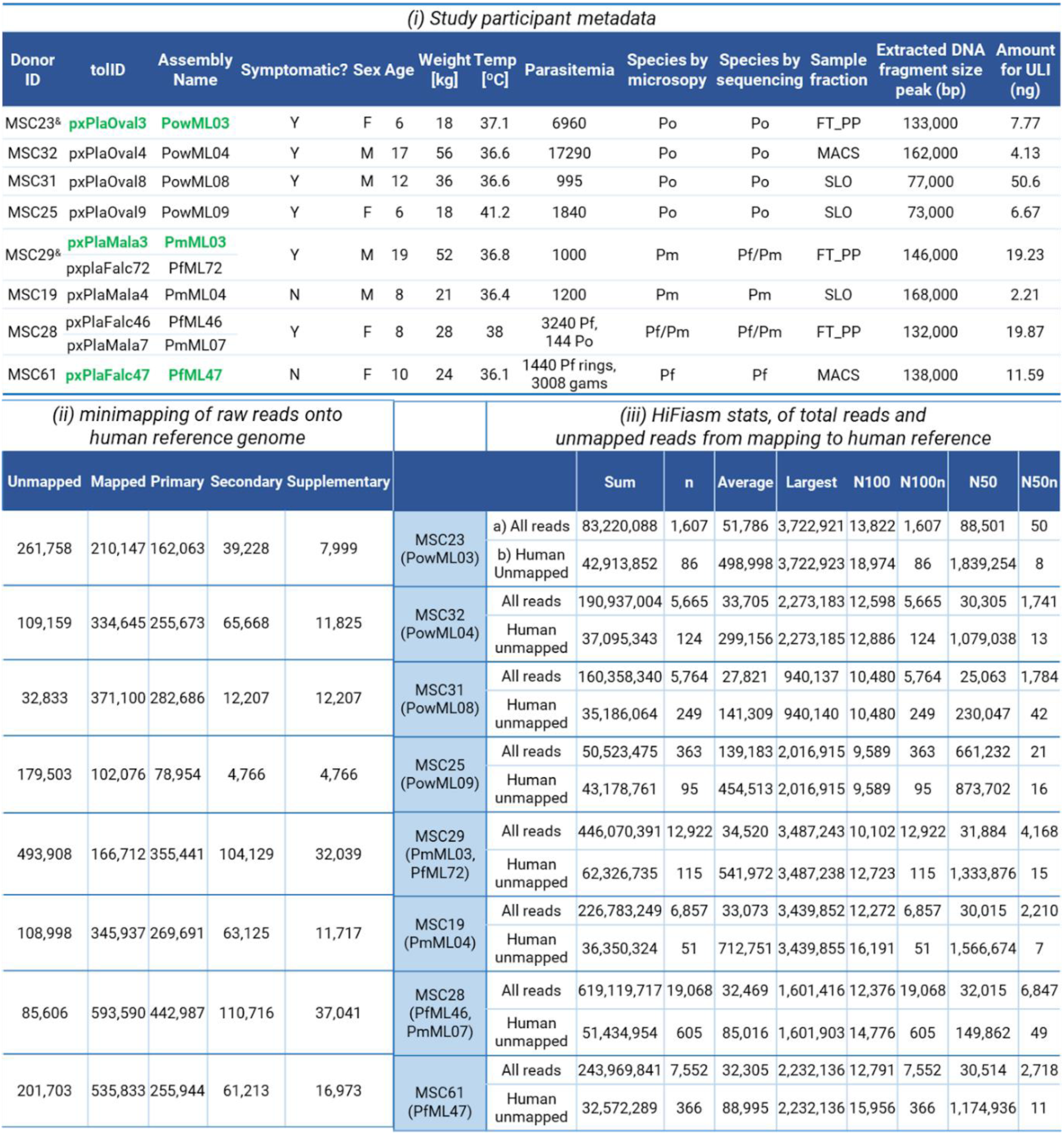
i, Metadata of study participants. Participants whose material is used for Hi-C are marked with a ‘&’. In bold green are the best quality assemblies for each of the species. Species are denoted as follows: Po - Plasmodium ovale spp., Pm - Plasmodium malariae, Pf - Plasmodium falciparum. Sample fractions used for data generation are denoted by MACS, SLO and FT_PP, described in detail in the Methods Section: MACS - MACS enriched fraction, SLO - Post-MACS, WBC-depleted blood lysed with Streptolysin-O, FT-PP - Post-MACS, WBC-depleted blood lysed with Saponin. Other notations are indicated as follows: Y - Yes, N - No, M - Male, F - Female. MSC25 was a P. falciparum/P. ovale spp. mixed infection sample, with P. falciparum. ii-iii, Mapping and assembly statistics for human read filtering and de novo assembly, summarizing the different stages of data processing. Mapping results against the human reference genome (ii), indicating the proportion of reads originating from the host, and Hifiasm assembly statistics (iii) generated using (a) all raw reads and (b) only reads that did not map to the human genome.

**Supplementary Table S2.**
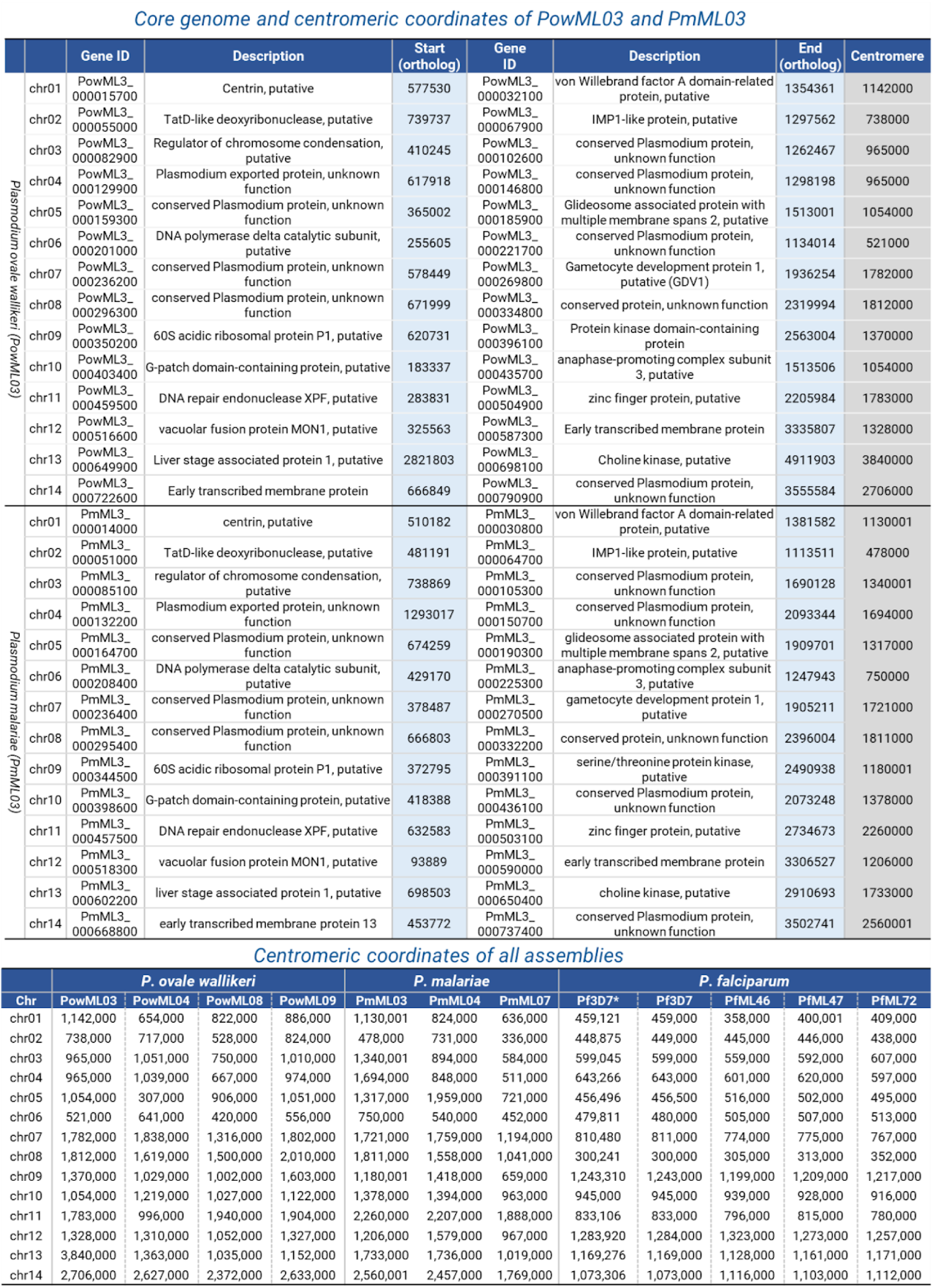
Core genome boundaries for P. ovale wallikeri and P. malariae assemblies were defined by the two outermost conserved 1:1 orthologs across human-infecting Plasmodium species on each chromosome, listed here. These boundaries coincide with reductions in GC content across in 1 kb windows. Centromeres were identified from Hi-C contact maps using PretextView using the cross-shaped interaction pattern for P. ovale wallikeri and P. malariae assemblies, as seen in Fig. 1 and Extended Data Fig. 1. For P. falciparum, the characteristic drop in GC content (which also coincides with the centromeres in P. ovale wallikeri and P. malariae) was used instead to infer the centromeric regions. The region so determined was concordant with the centromeric regions identified in Pf3D7, denoted with * (Bunnik et al. 2019).

**Extended Data Fig. 1:**
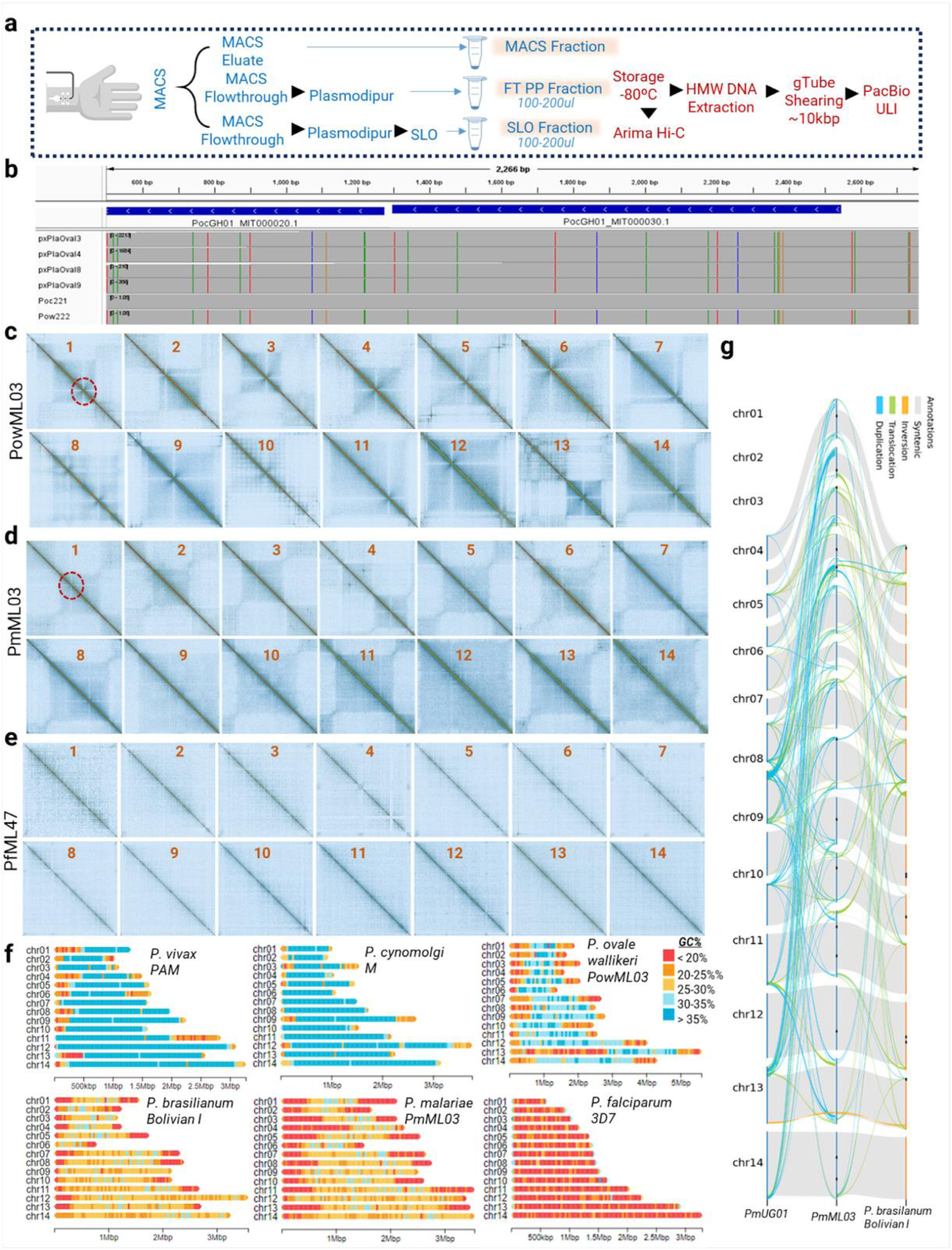
**a,** Sample workflow for enrichment of the parasite fractions directly from intravenous donor blood ^31,77^. The enriched fractions used for assembly and participants whose material is used for Hi-C are indicated with a ‘&’ in the Supplementary Table 1 showing the metadata of study participants. **b,** *cox1* gene locus from each of the *P. ovale* reference genomes sequenced here, compared to published *P. ovale curtisi* (PocGH01, PlasmoDB.v68 and Poc221, ^28^ and *wallikeri* reference genomes (Pow222, ^28^ confirm each genome is *P. ovale wallikeri*. Raw long reads of the *P. ovale* spp. specimens from this study and from Poc221 and Pow222 were mapped onto the *P. ovale curtisi* PocGH0 (PlasmoDB.v68) and visualised in IGV. Each vertical bar represents a SNP relative to PocGH01 and those from this study match the alt allele in Pow222. **c-e,** Pretext snapshots of individual chromosomes of (**c**) PowML03, (**d**) PmML03, and (**e**) PfML47. Cross-shaped interaction patterns typically associated with centromeric domains are indicated by circles, which were used to infer centromere coordinates, listed in **Supplementary Table 2**, of PowML03 and PmML03. Centromere coordinates of PfML47 were not ascertained this way as the stitch pattern is not apparent. **f,** Isochoric structure of the chromosomes of PowML03 is similar to species with similar GC content distribution, *P. vivax* PAM and *P. cynomolgi* M. Isochoric structure of the chromosomes of PmML03 is similar to species with similar GC content, *P. brasilianum* Bolivian I. *P. brasilianum*, a simian parasite was recently sequenced and assembled from *ex vivo* schizont material using high-fidelity PacBio long reads ^43^ and is regarded as nearly identical to *P. malariae*. *P. falciparum* doesn’t display a similar isochoric structure but does have high GC content in its telomeric ends (blue regions at the chromosome ends). **g,** SyRI synteny plots showing sequence similarity between the assemblies of PmUG01, PmML03, and *P. brasilianum* (Bolivian I).

**Extended Data Fig. 2:**
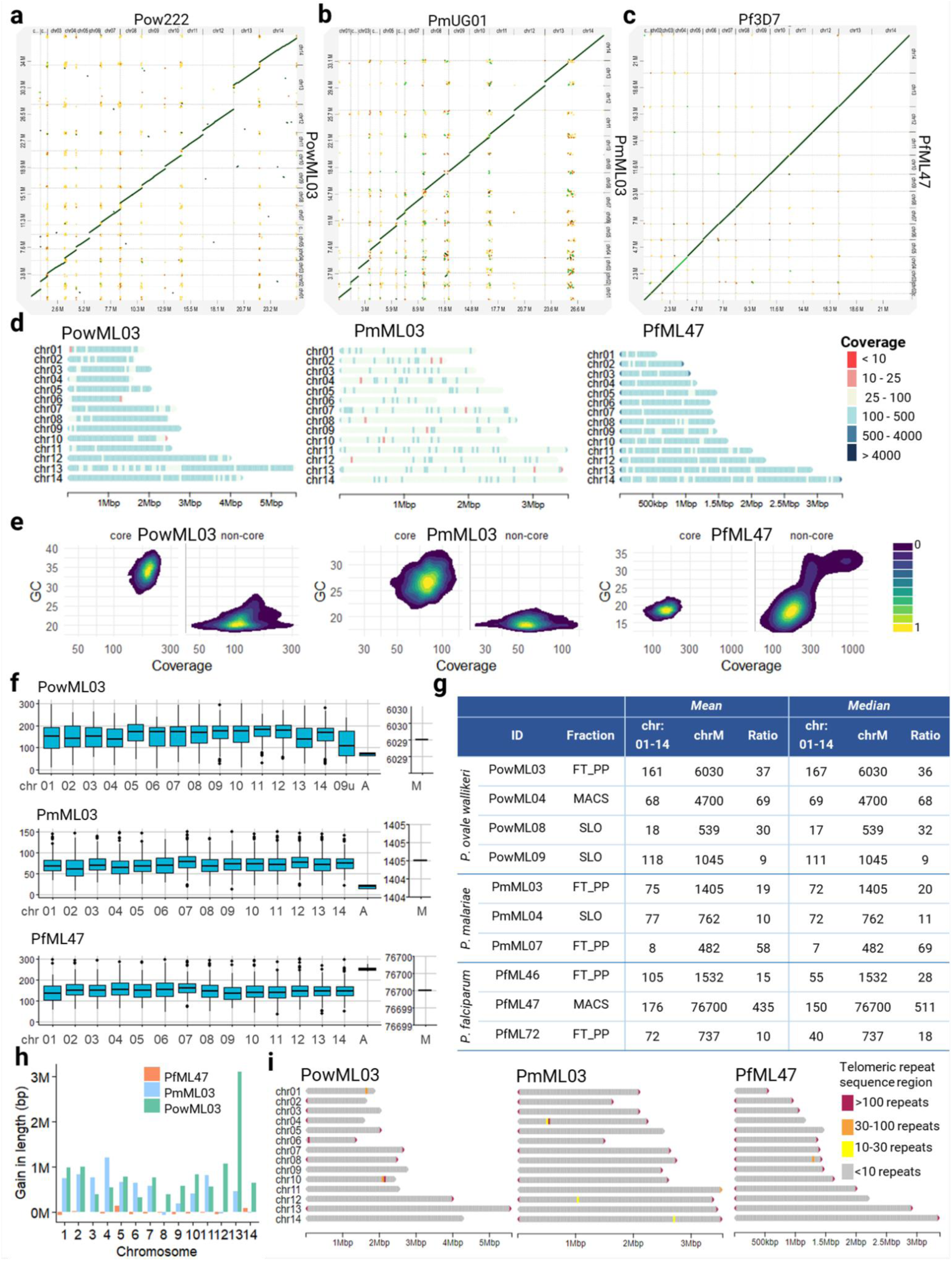
**a-c)** Chromosome dotplots of the best curated genome compared to the 14 chromosomes of current published reference genomes for (**a**) *P. ovale wallikeri* (PowML03) against Pow222 ^28^, (**b**) *P. malariae* (PmML03) against PmUG01 ^26^ and **(c)** *P. falciparum* (PfML47) against Pf3D7. Chromosomes in PowML03, PmML03 and PfML47 and the assemblies from other specimens of these species are ordered, oriented and named according to their syntenic relationship with Pow22, PmUG01 and Pf3D7, respectively. **d,** Coverage across each chromosome of the curated assembly for *P. ovale wallikeri, P. malariae*, and *P. falciparum*, binned into 10,000bp regions and colored according to the ranges displayed in the legend. **e,** Density plot showing coverage along with GC% of 10000 bp bins across all the chromosomes, split by their presence in the core or non-core regions, defined by the outermost position on each chromosome end of a 1:1 ortholog conserved across all human-infecting *Plasmodium* species. Extreme coverage regions are observed in the non-core regions of the chromosomes of PfML47 in particular. The plots show the density contours around the individual data points of each bin. **f,** Boxplots showing coverage across genomes of the curated assemblies of the specimens of *P. ovale wallikeri*, *P. malariae*, and *P. falciparum*, binned into 1000bp regions and displayed for each chromosome, apicoplast and mitochondrial contigs (in maroon, for chrM). **g,** Table showing the mean and median coverages for the curated assemblies across the nuclear genome and chrM, and fold change between them which can indicate the mitochondrial copy number. **h,** Gains in lengths of the chromosomes in the assemblies of *P. ovale wallikeri*, *P. malariae* and *P. falciparum* compared to their existing best references (Pow222, PmUG01 and Pf3D7). **i,** Telomeric repeat regions, inferred using telofinder, across the chromosomes of *P. ovale* PowML03, *P. malariae* PmML03, and *P. falciparum* PfML47.

**Extended Data Fig. 3.**
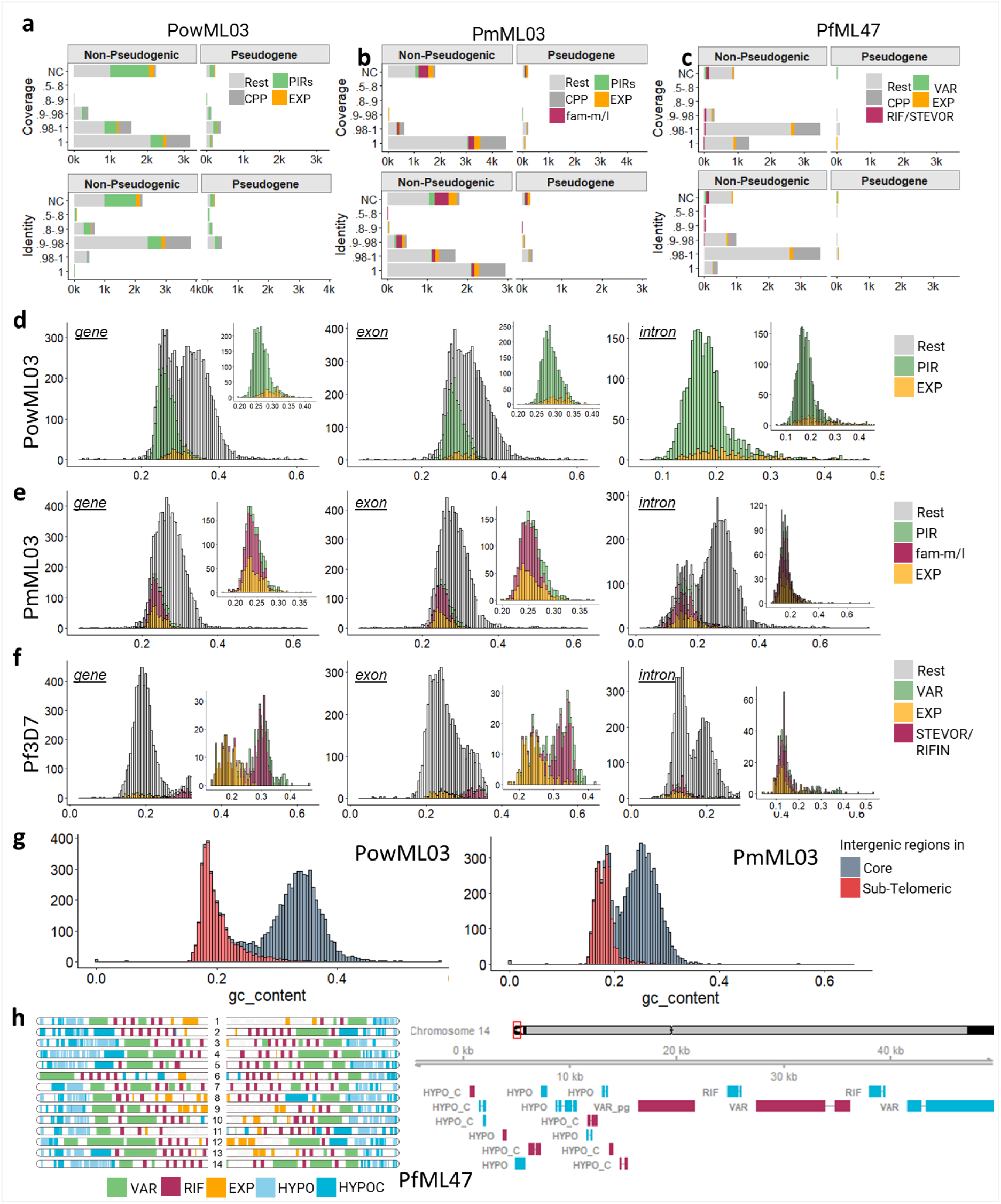
**a-c,** Identity and coverage of the transferred Companion annotations in **(a)** PowML03 **(b)** PmML03 and **(c)** PfML47. NC denotes non-Liftoff annotations (done by *ab initio* gene prediction by BRAKER2 and non-coding gene prediction by Aragorn), thereby lacking identity or coverage information relative to the source references used for annotations (PocGH01, PmUG01 and Pf3D7 respectively for PowML03, PmML03 and PfML47). **d-f,** Stacked histogram plots displaying GC% in the gene body, exonic, and intronic regions of genes belonging to the multigene families in (**d**) PowML03, (**e**) PmML03, and (**f**) Pf3D7, respectively, show their contribution to the skewed nucleotide ratio in the sub-telomeric regions. **g,** In addition to the gene body, intergenic regions are equally GC low in the sub-telomeric regions compared to the core genome. **h,** Companion annotations of “hypothetical proteins” beyond the *var* genes close to the telomeric ends are displayed across all the chromosomal ends of PfML47, along with an example of the gene distribution on the left arm of chr14. In the right panel, genes on the forward strand are colored blue and those on the reverse strand, maroon. Genes are grouped into the displayed gene families in **a-f and h** into categories as described in Methods.

**Extended Data Fig. 4:**
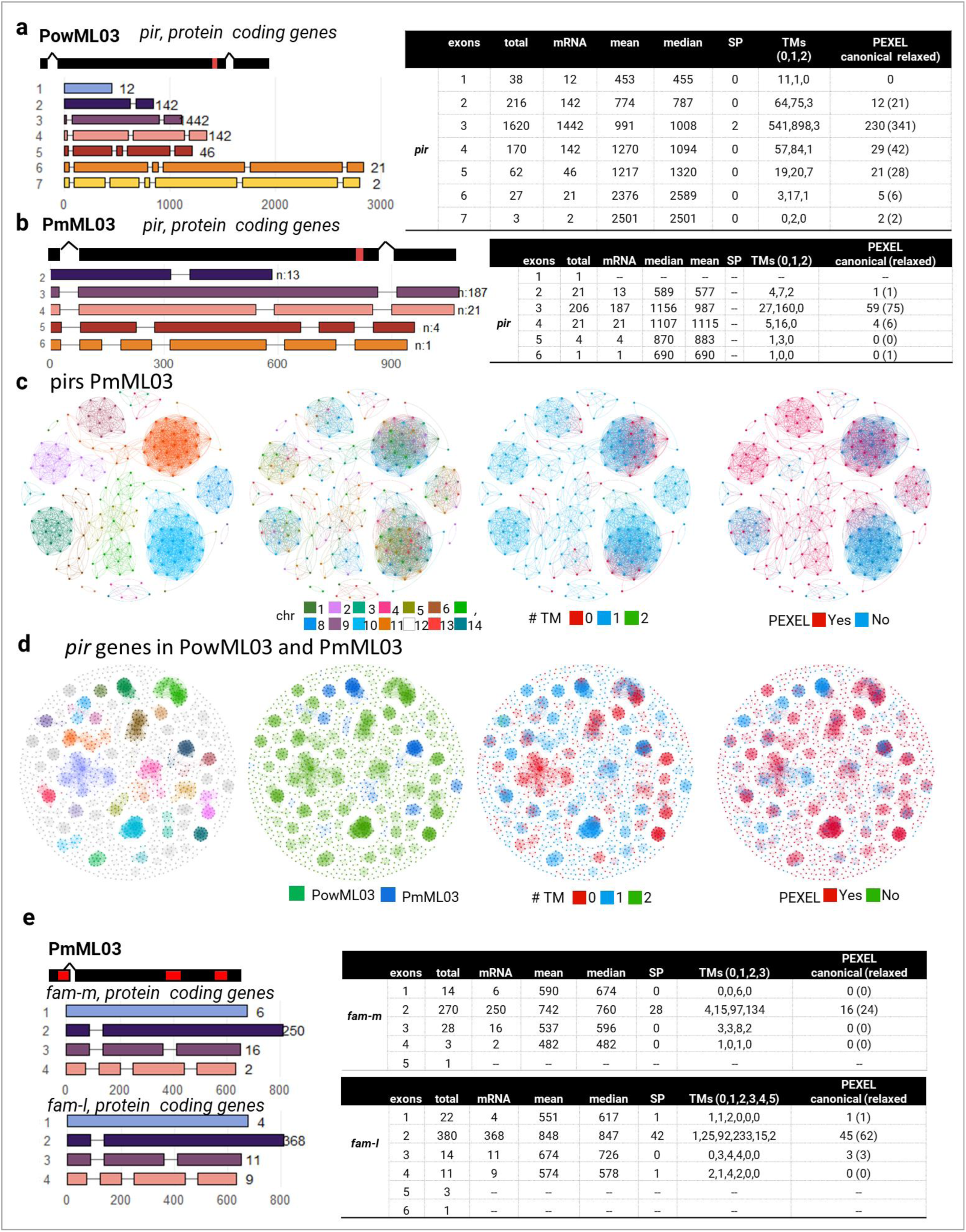
**a,** Exonic structures of PIRs in PowML03, displaying the most common example on top (black rectangles, with red block indicating transmembrane domain). **b,** Exonic structure of PIRs in PmML03, displaying the most common example on top (black rectangles, with red block indicating transmembrane domain). **c,** Protein similarity network of the *pir* genes in PmML03. Colored by module, by chromosome ID, by number of transmembrane domains (TM), and by number of exons. **d,** Protein similarity network of the *pir* genes in PmML03 and PowML03. Colored by module, species and by number of transmembrane domains (TM). Species form distinct, non-overlapping clusters, reflecting the sequence divergence of the pir gene family. **e,** Exonic structure of fam-m, fam-l genes in PmML03, displaying the most common example on top (black rectangles, with red block indicating transmembrane domain).

**Extended Data Fig. 5:**
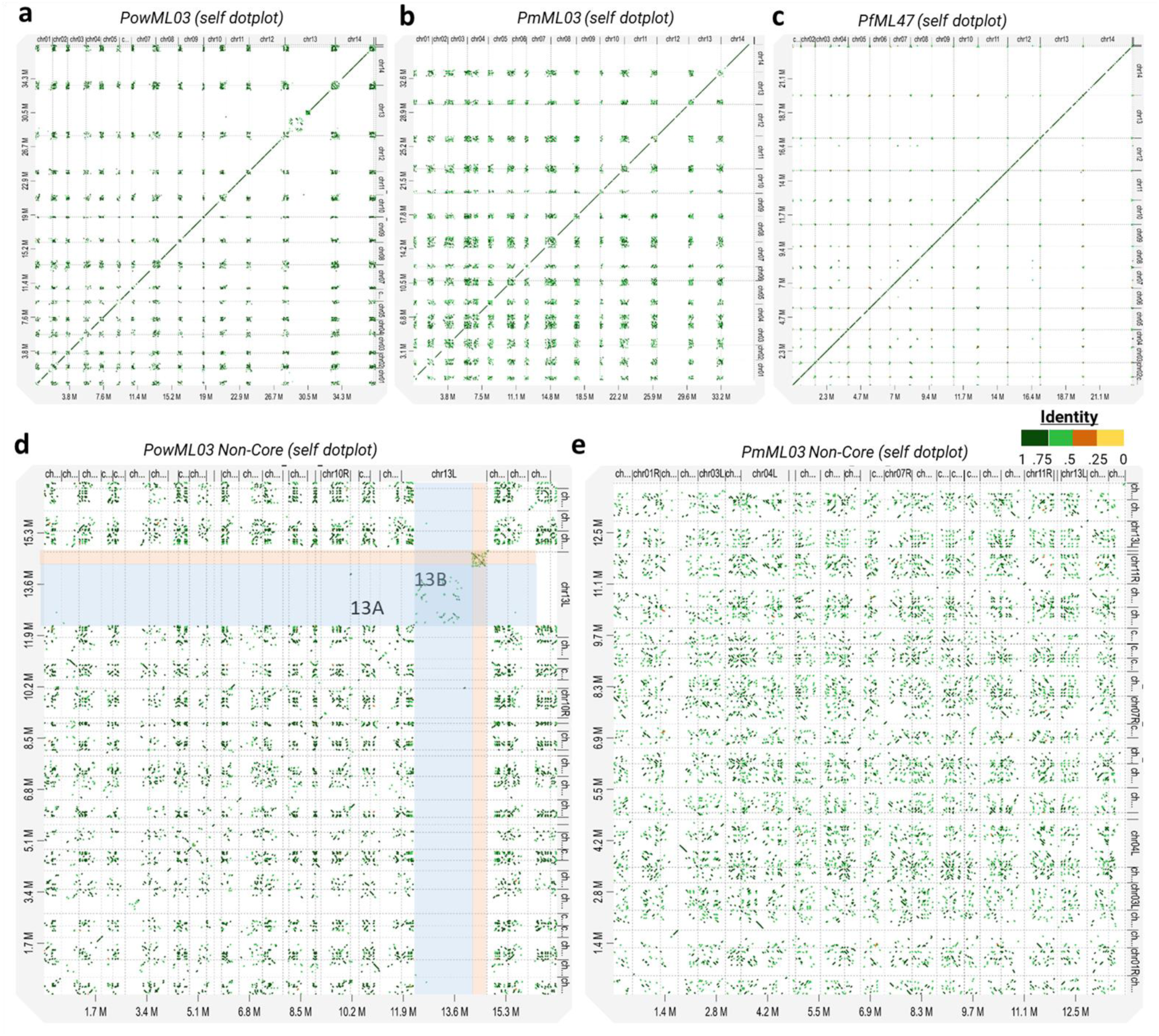
**a-c,** Self dot plot of assemblies of PowML03, PmML03 and PfML47, showing shared sequence identity across the subtelomeric regions. **d-e,** Self dot plot of only the non-core regions of PowML03 and PmML03. The dotplot in **(d)** highlights the left arm of chr13 which shares little similarity with the rest of the chromosomes. For all these dotplots, the paf (Pairwise mApping Format) file was generated by minimap2 with the options “-k15 -w10 -m40 --cs”, filtering out alignments less than 10,000 bp and plotted using D-genies ^35^.

**Extended Data Fig. 6.**
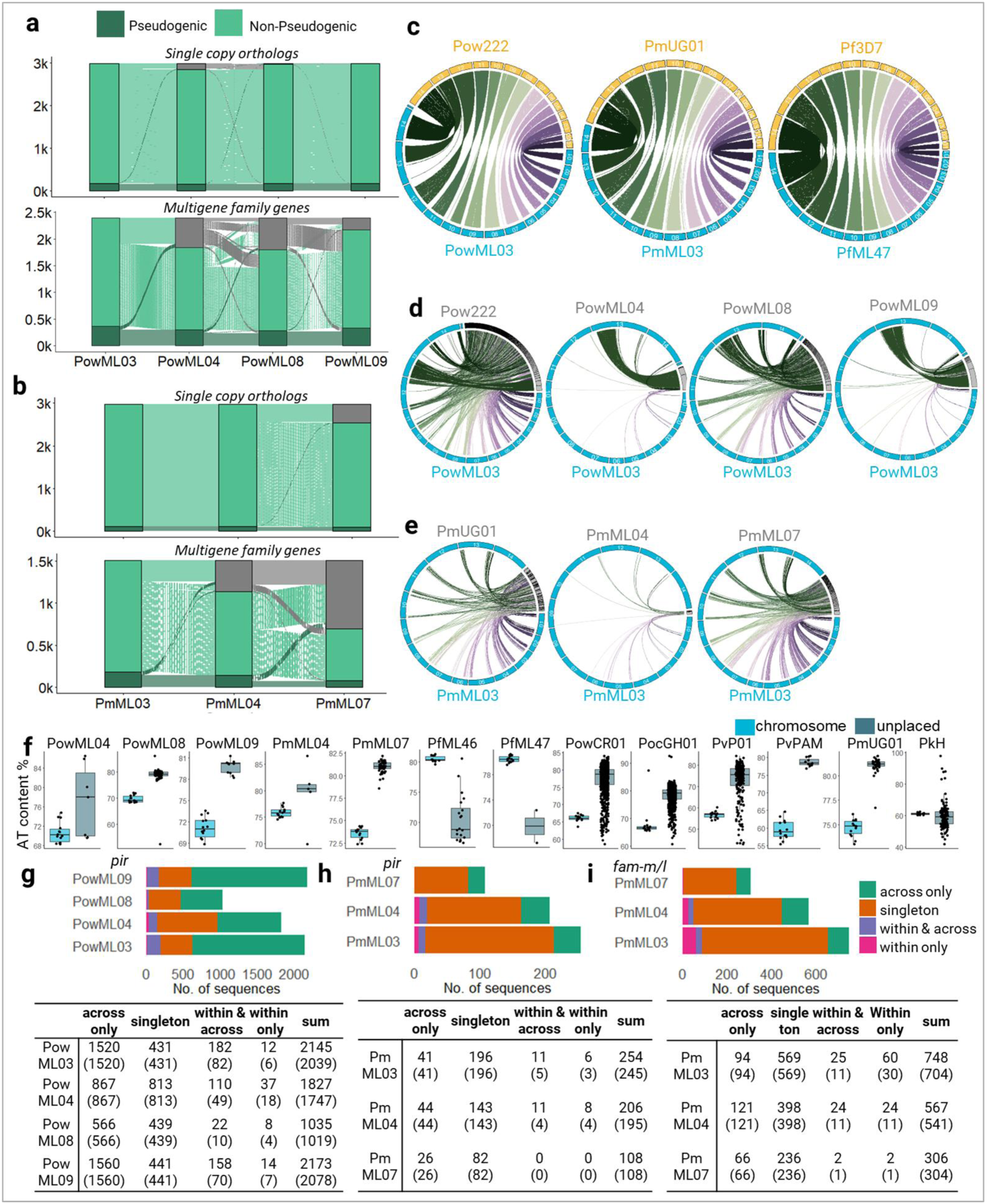
**a,** Annotations from PowML03 were transferred onto the other *P. ovale wallikeri* assemblies using Liftoff ^46^. Sankey plots were then used to evaluate gene transfer across specimens. While conserved single-copy orthologs within the core genome transferred reliably (top panel, small fraction of genes from PowML03, in grey, lacking its counterpart in PowML04, etc.), multigene families (bottom panel, ∼30% of genes from PowML03 lacking its counterpart in PowML04, etc.) in the subtelomeric regions showed poor transfer, likely reflecting the rearrangements and recombination characteristic of these regions. **b,** Same as **(a)** but for *P. malariae*. The relatively worse quality of the PmML07 assembly is reflected in the poor transfer of both conserved and multigene family annotations from PmML03 and PmML04. **c,** Using Liftoff-derived gene models within Companion, annotations from Pow222, PmUG01, and Pf3D7 mapped onto the PowML03, PmML03, and PfML47 assemblies, respectively, reveal clear patterns of conserved genomic regions, as visualised through Circos-based synteny plots. **d and e,** Feature based synteny of the unplaced contigs (grey) from lower quality *P. ovale* (**d**) (Pow222, PowML04, 08, 09) and (**e**) *P. malariae* (PmUG01, PmML04 and PmML07) assemblies was assessed in comparison to the best assembly generated here for each species. This shows that many unplaced contigs in poorer quality assemblies are found in the subtelomeric regions of the higher quality assemblies produced here. In particular, the two largest unplaced contigs in PowML04 and PowML09 are syntenic with the long sub-telomeric region of PowML03 chromosome 13. **f,** AT content of the unplaced contigs in the specimens from this study and references on PlasmoDB (v68). **g-i,** Stacked bar plots show the number of transcripts with sequences per dataset classified as unique singletons (orange), duplicated only within the same isolate (pink), only shared across multiple isolates (green), or present both within and across (purple) for **(g)** *pir* in PowML03,04,08,09, **(h)** *pir* in PmML03,04,07, and **(i)** *fam-m/l* in PmML03,04,07. Counts are shown in the table below for each category and sample, with the values in parentheses indicating the number of unique sequences amongst each category.

**Extended Data Fig. 7.**
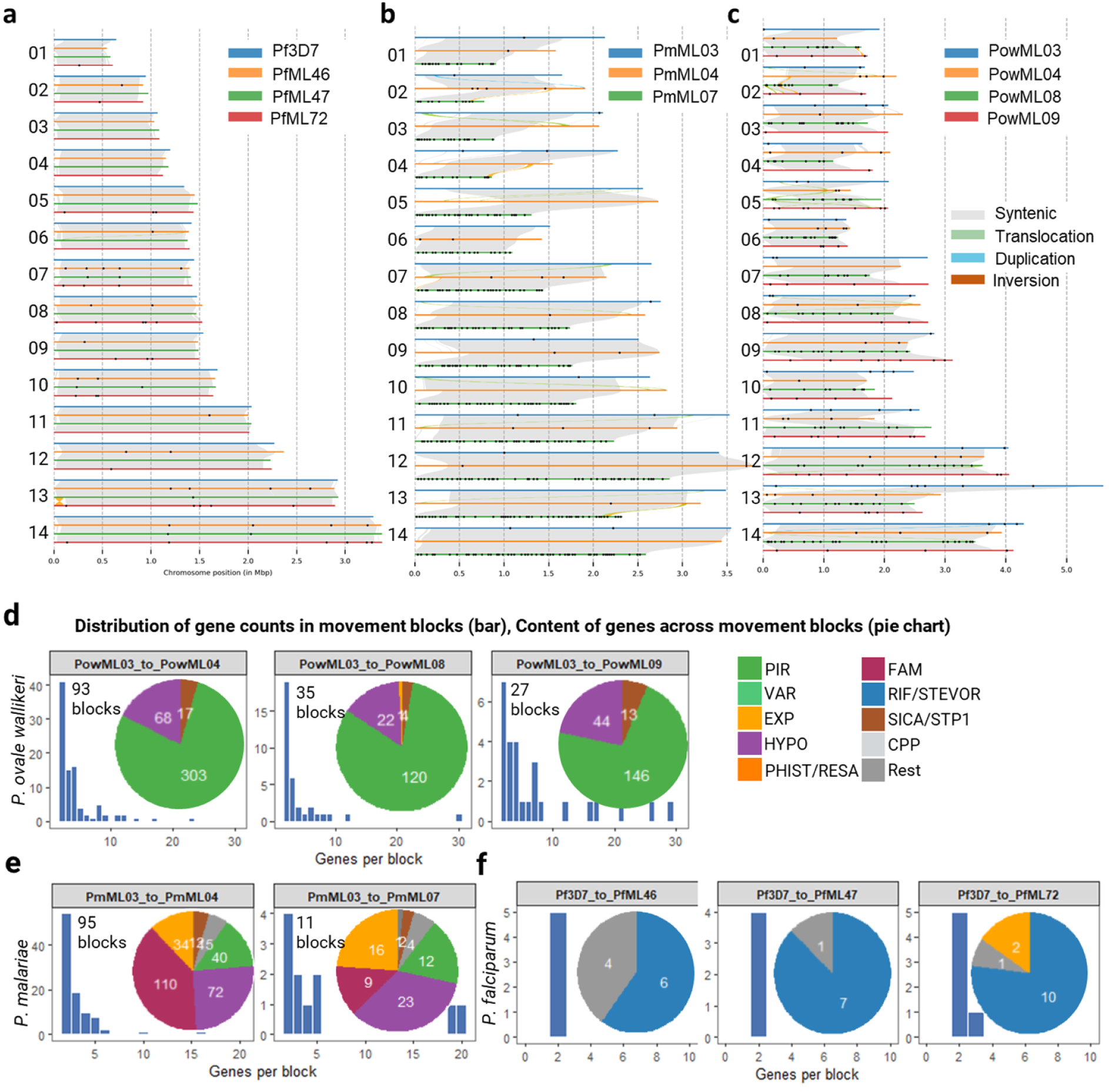
**a,** Synteny plots across the assemblies generated in this study for the three Plasmodium species showing synteny within the core genome, even across assembly breaks. For *P. falciparum*, Pf3D7 is included along with the other three specimens PfML46, 47 and 72. **d**, Gene movement blocks identified between *P. ovale wallikeri* assemblies (PowML03 with PowML04, PowML08, and PowML09)., **e,** Gene movement blocks between *P. malariae* assemblies (PmML03 with PmML04 and PmML07). **f,** Gene movement blocks between *P. falciparum* Pf3D7 and assemblies PfML46, PfML47 and PfML72. Bar plots show the distribution of gene counts per block, and accompanying pie charts summarise the composition of genes within blocks. The categories of gene families displayed in the table are described in Methods.

**Extended Data Fig. 8:**
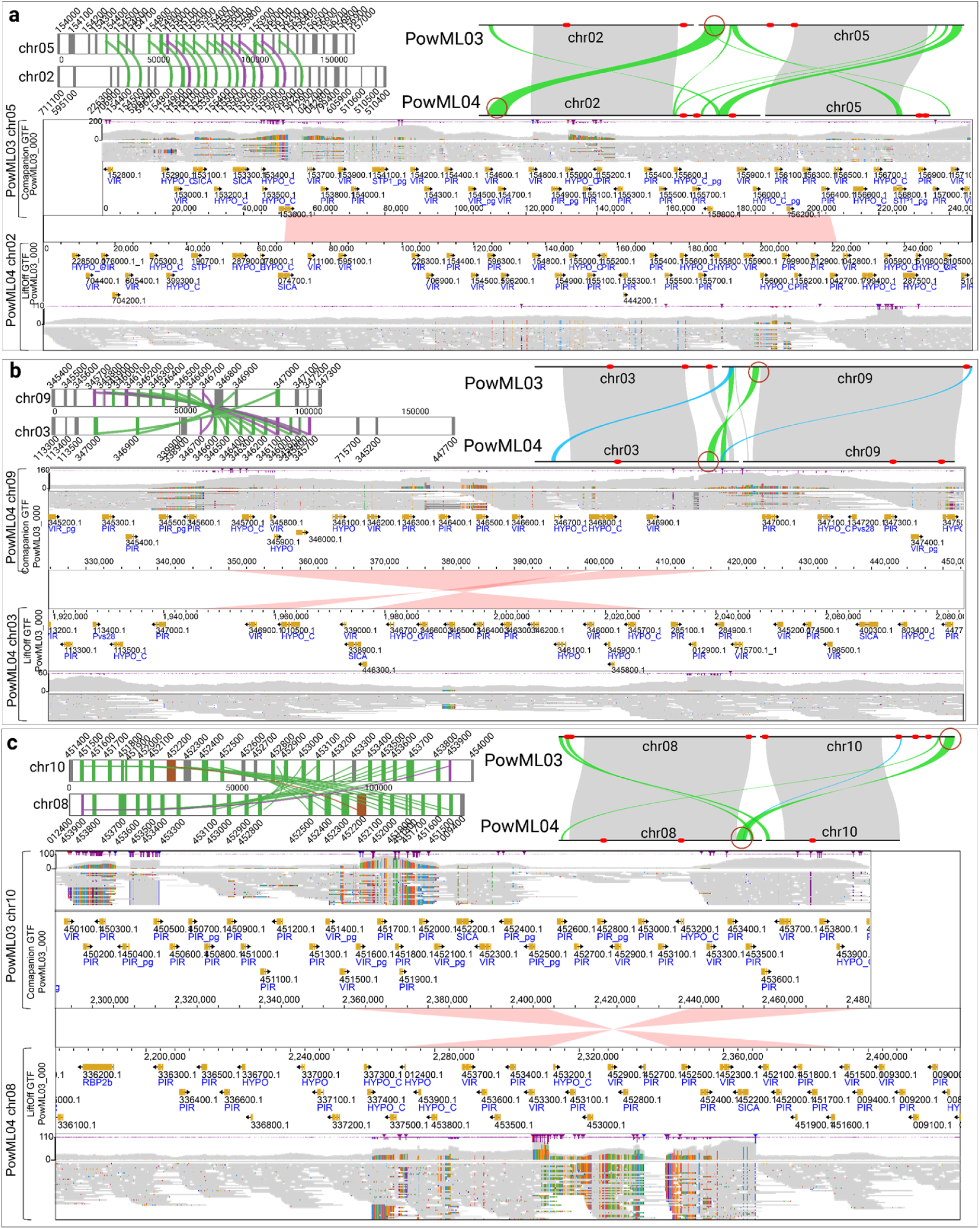
Examples of three loci in PowML03 identified as gene-block movements. For each example, the top left panel shows the syntenic block of genes, while the top right panel highlights the corresponding regions on the chromosomes using SyRI plots. The lower panels show JBrowse views of the corresponding loci in genomes PowML03 (above) and PowML04 (below). Gene annotations for PowML03 were generated using Companion. For PowML04, gene annotations were transferred from PowML03. **a**, A locus on chromosome 5 of PowML03 (above) shares sequence identity with a locus on chromosome 2 of PowML04 (below), as indicated by the shaded pink band connecting the two regions. The gene content within these regions show perfect concordance, whereas this concordance is lost beyond the boundaries of the syntenic block on either side. Raw reads mapping uniquely to each locus in their respective assemblies are shown alongside the gene annotations, with no evidence of breaks or inconsistencies, indicating that these regions could represent true genomic rearrangements rather than assembly artefacts. **b, c,** two further examples of similar dynamics.

**Extended Data Fig. 9:**
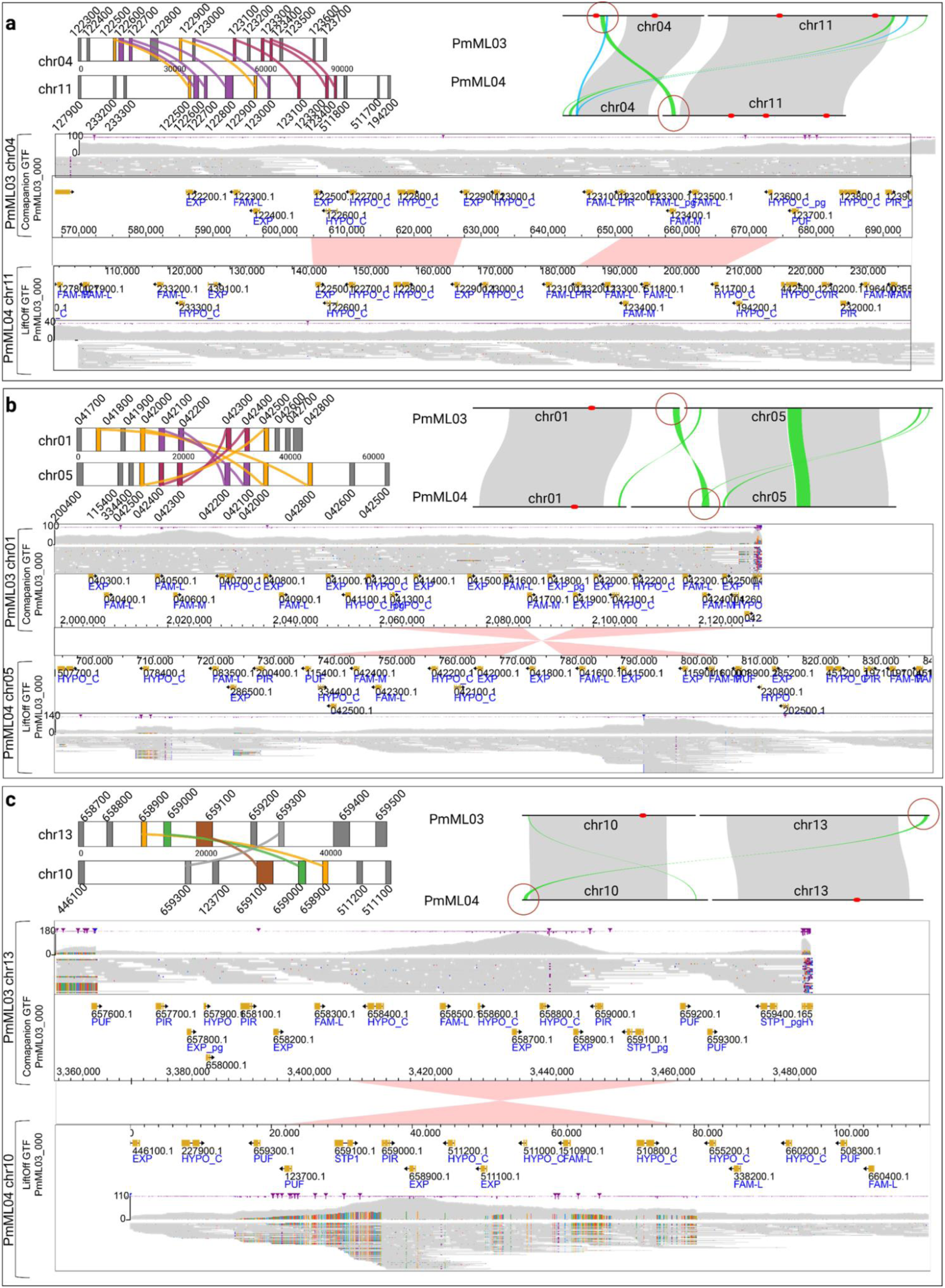
**a-c** same as EDF 8 but with three examples from *P. malariae*

**Extended Data Fig. 10:**
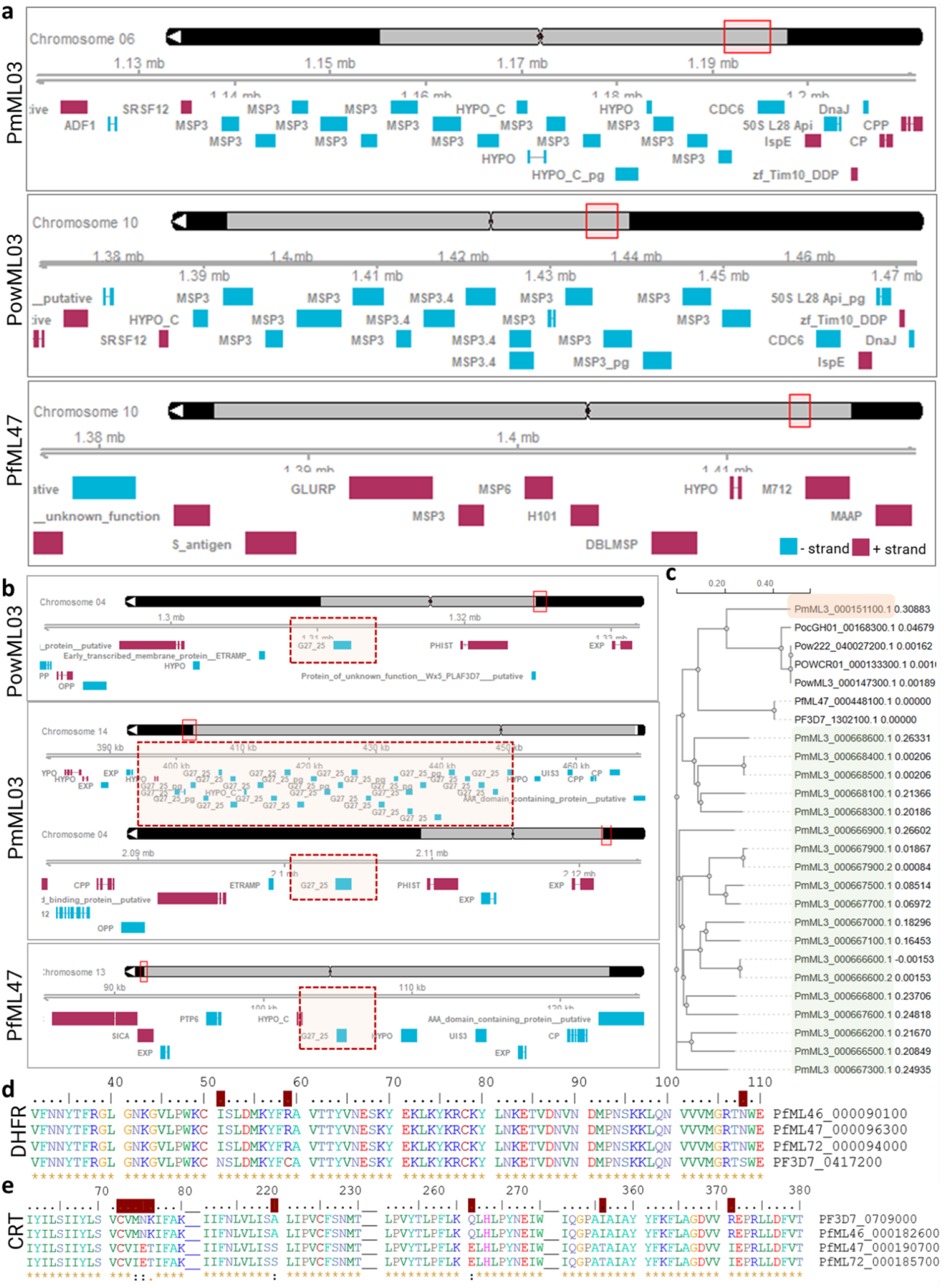
**a,** Genomic region surrounding *msp3* in *P. ovale wallikeri* PowML03, *P. malariae* PmML03, and *P. falciparum* PfML47. PowML03 and PmML03 show substantial expansion of the MSP3 locus, with 14 tandem copies on chromosome 10 in PowML03 and 13 tandem copies on chromosome 6 in PmML03. **b,** Genomic region surrounding *pfg27/25* in PowML03, PmML03, and PfML47, highlighting a marked copy number expansion in PmML03 on chromosome 14 and the single copy on chr04 (PmML3_000151100), in the middle panel. **c,** Phylogenetic analysis of Pfg25/27 protein sequences from the specimens shown shows that the expanded *pfg25/27* locus is phylogenetically distinct from the canonical locus on chr04 (PmML3_000151100), which instead clusters more closely with the single copy orthologs from other species. **d,** Alignment of DHFR sequences shows the triple mutant haplotype N51I/C59R/S108N, which is widely prevalent in Mali. The remainder of the sequence is identical to the 3D7 reference. **e**, Alignment of CRT sequences shows the CVIET mutant haplotype across residues 72-76, along with the A220S, Q271E, I356T and R371I mutations in PfML47 and PfML72. The remainder of the sequence is identical to the 3D7 reference.

